# Temporally and Functionally Distinct Contributions to Value Based Choice Along the Anterior-Posterior Dorsomedial Striatal Axis

**DOI:** 10.1101/2025.03.14.643367

**Authors:** Luigim Vargas Cifuentes, Edgar Díaz-Hernández, Myra M. Granato, Wenxin Tu, Kyuhyun Choi, Marc V. Fuccillo

## Abstract

While the dorsoventral and mediolateral organization of striatum has resolved clear functional distinctions, far less is known about how the anterior-posterior striatal axis contributes to behavioral control. We explore this within the dorsomedial striatum (DMS), a key region for value-based choice, by comparing population neuronal activity and function within anterior (A-DMS) and posterior (P-DMS) subregions while mice operantly seek reward. Neural recordings show that P-DMS encoded action values and strategy information prior to choice selection while A-DMS activity represented recently selected choices and their anticipated values via a dynamic population reorganization immediately following action selection. Optogenetic perturbations were consistent with these temporally distinct coding properties as unilateral manipulation of the P-DMS prior to choice biased choice contralaterally in a value-dependent manner and unilateral inhibition of the A-DMS following choice impaired future value-based action selection. Using anterograde tracing, we found that the A-DMS and P-DMS projected to a common region within the ventromedial substantia nigra pars reticulata (vmSNr), which contained value-related signals combining aspects of upstream DMS processing. Together, our results support a model for temporally distributed influence on value-based choice across the anterior-posterior axis of the DMS.

## INTRODUCTION

As the input nucleus of the basal ganglia, the striatum receives excitatory inputs from most forebrain projections including cortex, thalamus, amygdala and hippocampus^1–3^. As such, it is likely that the differential function of striatal territories is strongly determined by distinct afferent inputs processed by a canonical striatal microcircuit featuring two SPN subtypes and sparse local interneurons^4,5^. The dorsal-ventral (DV) and medial-lateral (ML) striatal axes have been extensively characterized with functionally distinct contributions to behavior aligning to their predominant inputs^6^ – the ventral striatum receives limbic afferents and processes reward approach; the dorsomedial receives associative afferents and mediates goal-based action selection; the dorsolateral striatum receives sensorimotor inputs and has functions in motor output and habitual responding. Although less studied, converging lines of evidence suggest a unique functional architecture of the striatum along its anterior/posterior (A/P) axis. Early tracing studies in NHPs and rats consistently highlighted the existence of longitudinal projection patterns for cortical neurons synapsing in striatum^7–9^. Recent studies have emphasized that the rostro-caudal origin of cortical inputs does not directly track with their A/P target within striatum – in one example, the caudal striatal tail is strongly innervated by diametrically opposed prefrontal cortical regions^10^; in another, simultaneous retrograde tracing from distinct parts of the DMS A/P axis labeled intermingled but distinct populations in prefrontal and sensori-motor cortical areas^11^. These discoveries highlight a notable gap in our understanding of striatal function along the anterior-posterior (A/P) axis^12–14^.

Goal directed behaviors are a central adaptive behavior which allow organisms to select strategies and motor outputs that optimize valued outcomes such as obtaining reward or avoiding exposure to aversive events. Successful execution of these behaviors requires filtering of outcome-relevant environmental cues, selection of appropriate actions, evaluating the consequences of those actions and using that information to guide future choice. The neural ‘binding’ of temporally displaced outcomes and their preceding actions (‘credit-assignment’) remains a central unanswered question. Prior studies have highlighted the importance of DMS, the rodent analogue of the primate caudate nucleus, for goal-directed behavior via both functional manipulations and discovery of neural representations of relevant decision variables. Early excitotoxic lesion studies suggested that the posterior regions of the DMS (termed ‘P-DMS’) but not more anterior parts (‘A-DMS’) could support outcome devaluation-mediated reductions in responding^15–17^. While multiple other studies have confirmed the role of P-DMS in instrumental performance according to action-outcome contingencies^18,19^, the role of the A-DMS has been less defined, with studies uncovering functions in supporting reversal learning, win-stay/lose-shift strategies and stimulus-outcome associative learning^11,17,20,21^. Our understanding of functional divisions of DMS along the A/P axis remains limited by lack of studies that have systematically investigated this organizational framework by performing recordings and perturbations across DMS sub-regions within the same behavior.

Another understudied aspect of regionally distributed striatal processing is the mechanisms by which functionally related computations are integrated to produce holistic control of motor output. Most neural signals entering the rodent striatum make their way to the largest basal ganglia output nuclei, the substantia nigra, pars reticulata (SNr)^22^. Careful anatomical tracing studies have demonstrated that there is a topographical organization of striatal outputs to the SNr and subsequently from the SNr to downstream thalamic and collicular nuclei^23^. These circuits are thought to be essential for behavioral orienting, attentional control as well as saccade and motor release. While these pathways form segregated closed-loop circuits according to their D-V and M-L striatal origin^23^, the potential for overlapping target field integration from A-P striatal regions remains unknown. Recordings in SNr have identified both signals for movement and reward as animals engage in a goal-directed tasks^24–26^. Interestingly, in a visual object value-association task, stable rather than flexible value associations were selectively represented by SNr neurons^26^. Furthermore, the rate and pattern of SNr spiking correlated with efficient value-based scanning of visual objects^27^. Despite these interesting advances, our current knowledge of how behaviorally relevant striatal signals are propagated to the SNr during value-based choice remains limited.

In the present study, we sought to examine functional differences of basal ganglia circuits along the A/P axis by using *in-vivo* recording and optogenetic techniques while mice performed a volume-comparison task. *In vivo* extracellular recordings of neurons within the A- and P-DMS demonstrated divergent encoding of behavioral events, choice strategy, and subjective value with distinct temporal dynamics. Specifically, we found that P-DMS population modulation began at the beginning of each trial, prior to choice while the largest reorganization of A-DMS populations occurred immediately after choice selection, persisting throughout the trial remainder. Consistent with this distributed temporal processing, we found that manipulations in the P-DMS were only effective in the pre-choice epoch, and they induced a value-dependent bias towards the contralateral side. In contrast, perturbations in the A-DMS were only effective during the post-choice period, altering the animals’ use of outcomes on future choice. Using viral-based anterograde tracing techniques to analyze the outputs of A- and P-DMS in SNr, we found that, despite broad segregation, the ventromedial SNr (vmSNr) was a potential site of convergence between these loops. Extracellular recordings targeted to this region showed neural encoding of value-based choice that resembled aspects of both A- and P-DMS. Together, our data suggest a new model of temporally and functionally distinct control of decision-making across the A/P axis of the DMS that is subsequently integrated within the SNr.

## RESULTS

### A task probing distinct value-based choice contexts

To systematically investigate functional differences between the A-DMS and P-DMS, we used a volume-based choice task^28^ wherein mice relied on outcome feedback to select the better of two options. Mice were trained in a freely moving volume-comparison task with fixed block lengths of 25 trials and uncued fluctuating optimal sides, defined by larger reward volume (Fig.1A,B). Mice initiated trials via center port poke, registered choice in the two lateral ‘choice’ ports and returned to the center port to ‘reveal’ the outcome resulting from their prior choice. We varied the difficulty of the value comparison by opposing the 12µl high value side with either a small volume alternative (4µl, 12v4 block) or no reward (0µl, 12v0 block) in random block-wise fashion (Fig.1B). To increase choice sampling, we omitted reward on 30% of trials, applied to both choice ports (labeled as ‘noise’ in Fig.1A). Finally, to promote unbiased choice selection at block start, each block concluded with 5 non-rewarded trials (labeled hereafter as “extinction”). Overall, mice performed nearly 500 trials with this structure per hour-long session.

**Figure 1.**
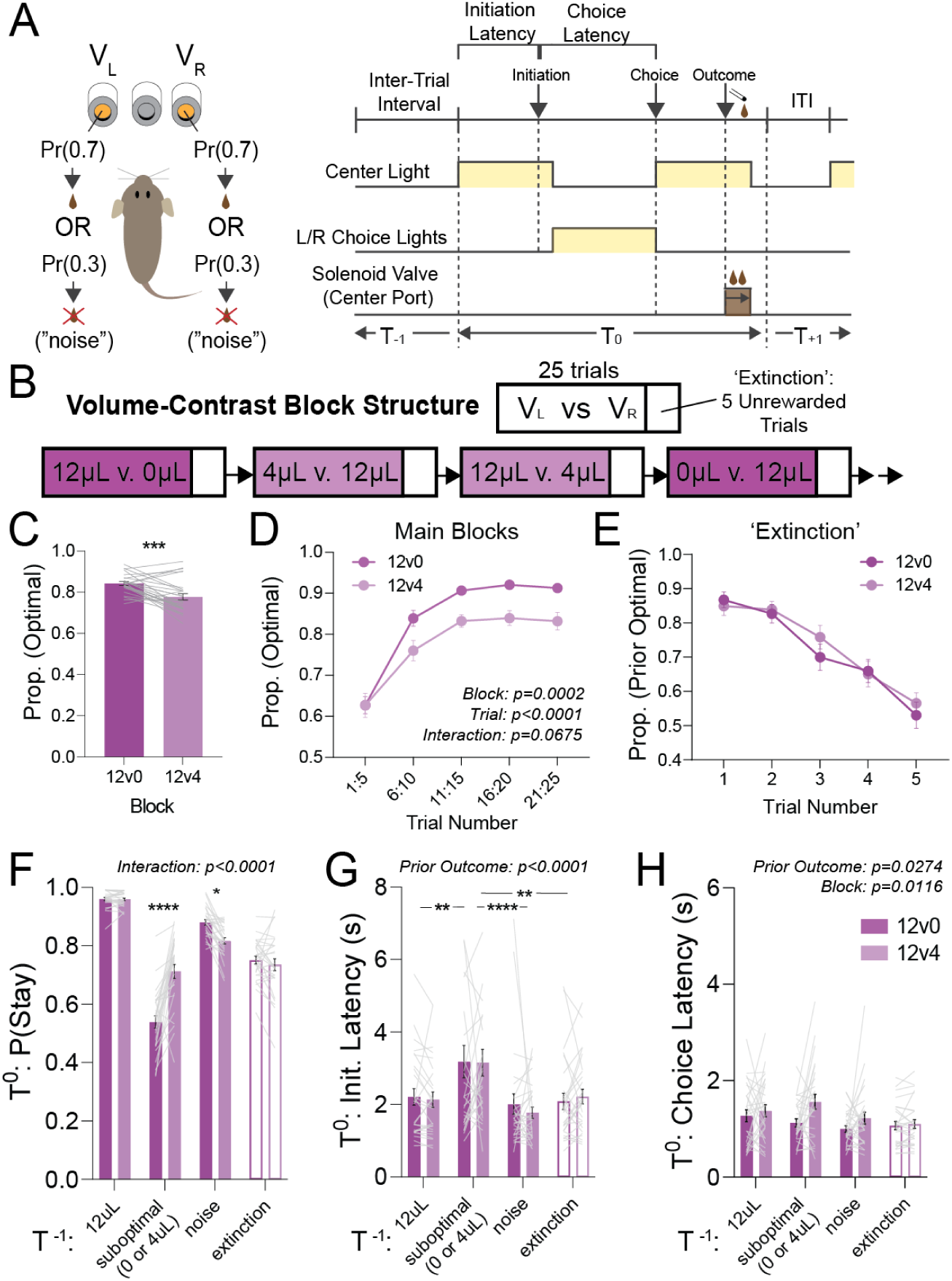
A volume contrast task evokes distinct behavioral performance. (A) Self-paced two-alternative choice task. Animals initiate a trial by poking into the center port, then make a choice by choosing the left or right port. Reward is collected in the center port. Each choice has a 70% probability of giving the current block reward outcome and a 30% probability of delivering no reward regardless of choice (‘noise’ trial). (B) Block structure. Each block is fixed at 25 trials in length with one side yielding 12µL and the other side yielding either 4µL or 0µL. Between every block, there are 5 unrewarded trials that probe the outcome-sensitivity of choice (labeled ‘extinction’). The 12µL side switches between each block. (C) Animals make fewer optimal choices in the 12v4 blocks compared to 12v0 blocks (*n*=27; Accuracy: ***P=0.0002, Wilcoxon matched pairs rank sum test). (D) Average trial accuracy binned in groups of five trials is lower across 12v4 blocks compared to 12v0 blocks (*n*=27; RM-two-way ANOVA, trial: F_2,55_ = 70.80, ****P < 0.0001, block: F_1,26_ = 19.50, ***P = 0.0002, interaction: F_3,67_ = 2.600, P = 0.0675). (E) Average responding during unrewarded ‘extinction’ sections is not different following 12v0 blocks or 12v4 blocks (*n*=27; RM-two-way ANOVA, trial: F_3,82_ = 50.06, P < 0.0001, block: F_1,26_ = 0.3234, P = 0.5744, interaction: F_3,72_ = 0.6567, P = 0.5705). (F) Average probability of returning to prior choice on current trial varies by prior outcome. Animals are more likely to return following 4uL rewards on the suboptimal side compared to 0uL. Animals also respond to noise differently between blocks. (*n*=27; RM-two-way ANOVA with Šídák *post-hoc* multiple pairwise comparison, interaction: F_3,208_ = 22.95, ****P < 0.0001, prior outcome: F_3,208_ = 174.8, ****P < 0.0001, block: F_1,208_ = 4.738, *P = 0.0306). (G) Average initiation latency following suboptimal reward outcomes is higher than latencies following 12uL or noise outcomes (*n*=27; RM-two-way ANOVA, interaction: F_3,208_ = 0.1461, P = 0.9321, prior outcome: F_3,208_ = 8.038, ****P < 0.0001, block: F_1,208_ = 0.07284, P = 0.7875). (H) Average choice latency is higher in 12v4 blocks compared to 12v0 blocks and differs according to the previous outcome (*n*=27; RM-two-way ANOVA, interaction: F_3,208_ = 1.274, P = 0.2845, prior outcome: F_3,208_ = 3.110, *P = 0.0274, block: F_1,208_ = 6.479, P = 0.0116).

We found that mice exhibited different behavior in the 12v0 and 12v4 blocks, measured along multiple parameters. Mice had higher overall accuracy in the 12v0 blocks compared to the 12v4 blocks (Fig.1C), with these patterns emerging after the first 5 trials of a new block (Fig.1D). In contrast, mice returned to near chance responding after 5 extinction trials (neither response produced reward) regardless of block identity (Fig.1E). To explain the behavioral differences in 12v0 and 12v4 blocks, we examined how animals responded to each type of task feedback. We found that animals returned to large volume ports with a high probability in both block types (Fig.1F). In 12v0 blocks, animals returned to the small volume side (in this case 0ul) ∼50% of the time while in 12v4 blocks, animals returned to lower value side ∼70% (Fig.1F). Choice efficiency can also be impacted by how animals respond following the 30% unrewarded noise condition. We found that animals in the 12v0 blocks were better able to ignore noise and remain on the high value side as compared to 12v4 blocks (Fig.1F). Thus, the lower accuracy in 12v4 blocks is due to the increased difficulty of comparing different reward volumes and tracking the high-value side following noise trials. To examine block-specific differences in behavioral execution, we analyzed task latencies with respect to prior and current trial outcome. Consistent with previous results, mice were slower to initiate subsequent trials following suboptimal outcomes in both block types (Fig.1G). In contrast, latency to choice following all outcomes was faster in 12v0 blocks and slower in 12v4 (Fig.1H). Overall, performance during ‘extinction’ trials was distinct from performance in the rewarded parts of 12v0 and 12v4 blocks. Animals were more likely to persist in choosing the previously rewarded side in extinction than following suboptimal outcomes in the main blocks (Fig. 1F). Furthermore, animals did not exhibit an increase in initiation and choice latency following extinction trials, as seen with suboptimal outcomes in the main blocks (Fig. 1G-H). These data suggest that extinction trials are characterized by reduced outcome sensitivity as compared to trials within 12v0 and 12v4 blocks. Finally, we used logistic regression to examine the impact of choice and outcome history on current trial choices (Fig.S1A-D). We found that the prior three rewarded trials all contributed to current choice, suggesting multi-trial integration of reward history. Taken together, these data show that animal choice is sensitive to outcome value and that 12v0 and 12v4 blocks evoke distinct strategies, choice patterns, and motor performance.

### Differential recruitment of the A-DMS and P-DMS as animals perform the volume-comparison task

Several studies have shown that the A-DMS and P-DMS differentially contribute to action-outcome learning^15,16,19^. Other work has recorded DMS spiny projection neurons (SPNs) during value-based tasks but didn’t account for their location along the A-P axis^29,30^. Finally, recent work has shown that neuronal encoding and function of prefrontal inputs for value-based choice are segregated by their target sites along the anterior-posterior extent of DMS^11^. To systematically investigate whether DMS sub-regions exhibited differences in the processing of value-based decisions, we placed movable electrodes in either A-DMS (A/P: +1.3→ 0.6) or P-DMS (A/P: 0→ -0.4) while mice performed our volume-based choice task (Fig.2A,B). We recorded 148 isolated units in the A-DMS (N=5 mice) and 112 units in the P-DMS (N=6 mice; see Methods for full details). We first stretched individual trial firing rates to global trial averages to remove trial-by-trial temporal variability (hereafter labeled ‘warped’; Fig.2C,D). To broadly compare overall population dynamics between A- and P-DMS, we performed principal component analysis on each isolated neuron’s average firing rates. The first three principal components of each population (accounting for 54.1 and 47.8% of firing rate variance for A- and P-DMS, respectively) exhibited distinct trajectories – the A-DMS exhibited elevated activity during the ITI followed by suppressed firing beginning at task initiation and continuing until outcome presentation (Fig.2E). In contrast, the P-DMS exhibited coordinated periods of increased activity during the ITI, preceding choice, and immediately after choice (Fig.2F). Consistent with this, 3D mapping of the first 3 principal components exhibited longer trajectories during the post-choice and ITI for A-DMS and the initiation-choice period for P-DMS (Fig.2G,H).

**Figure 2.**
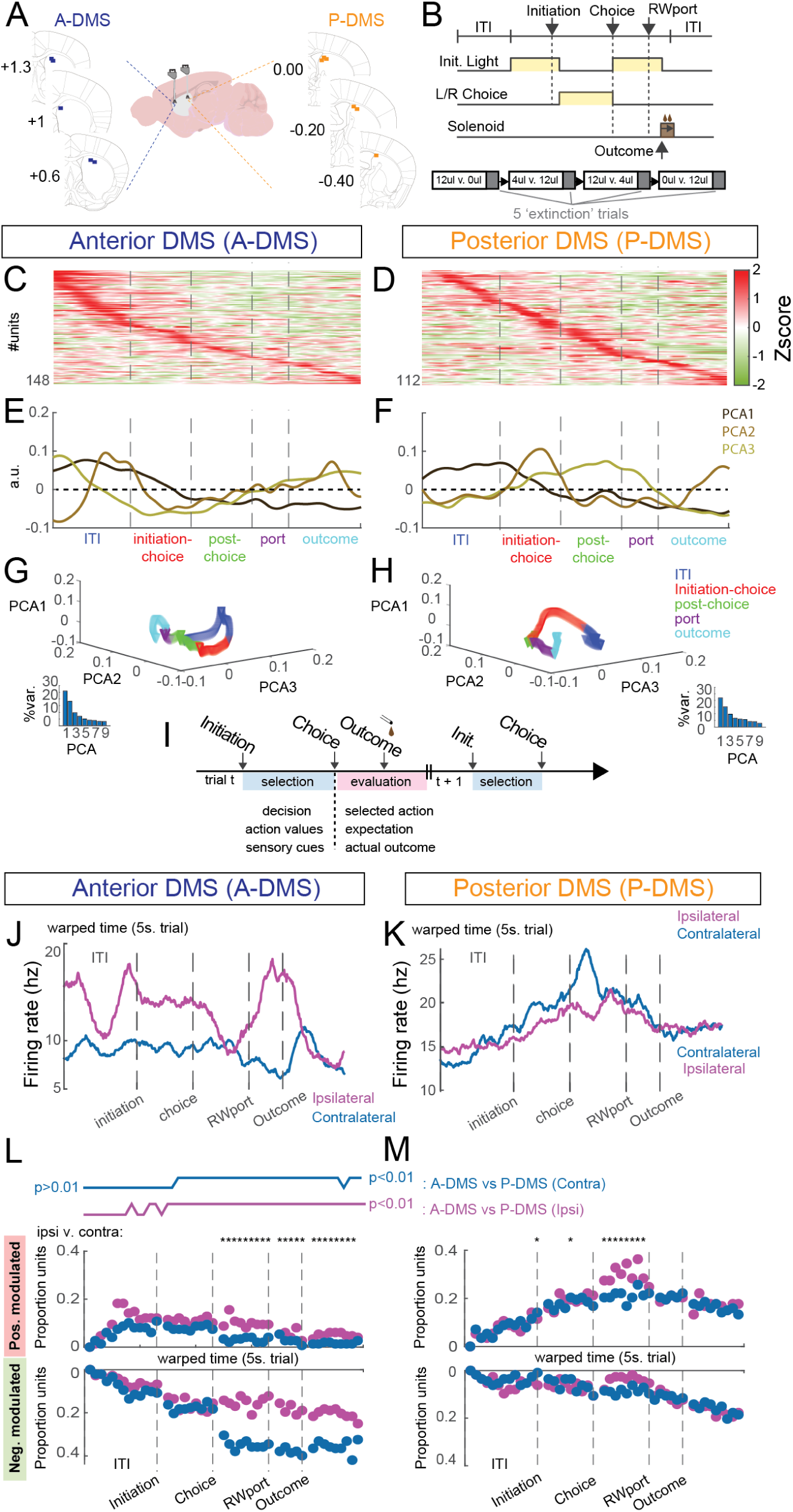
The A- and P-DMS exhibit broadly divergent patterns of modulation. (A) Anatomical localization of the movable recording electrodes (start site) placed in either A-DMS (n=5; A/P: +1.3→ +0.6) or P-DMS (*n*=6; A/P: 0→ -0.4) (B) Structure of the volume contrast task. (C and D) Averaged activity of A-DMS (C) and P-DMS (D) units recorded in the volume block task (time warped) aligned to trial start. Units are sorted by peak activity. (E and F) The first three principal components of the averaged firing rate in A-DMS (E) and P-DMS (F) demonstrate different global patterns of task-aligned activity. (G and H) Trajectories for the averaged trial-aligned firing rate across the first three principal components show longer trajectories in the post-choice epoch for the A-DMS (G) and in the initiation-choice epoch for the P-DMS (H). Bottom panels show the variance explained by successive principal components. (I) Schematic detailing task-relevant behavioral variables and their temporal location within the task. (J and K) Example individual striatal neuron firing rates that differ with respect to whether choices are ipsilateral or contralateral to the recording site in A-DMS (J) or P-DMS (K). (L and M) Top: Comparison of A-DMS and P-DMS population modulation showing differences for contralateral (blue line) and ipsilateral choice (magenta line) beginning largely at trial initiation (Fisher exact test p<0.01). (L) Proportion of positively (red) and negatively (green) modulated A-DMS units across the trial shows ipsilateral vs. contralateral differences beginning after choice and lasting until trial conclusion (* indicates p<0.0001 using McNemar’s test corrected for multiple comparisons). (M) P-DMS modulated units diverge for population encoding of choice briefly following choice selection.

To define the task-specificity of neural signals, we compared single unit activity in each striatal region during the first 25 trials, where mice used feedback to optimize choice, to the last five trials, where neither choice was rewarded (designated as ‘extinction’; Fig.2B). We used sliding-window receiver operating characteristic (ROC) analysis to compare activity in sequential 500ms time bins (advanced in 50ms increments) against ITI baselines for trials 1-25 versus trials 26-30. Within every time window, each unit’s auROC value was thresholded into a modulation score – either positive, negative, or non-modulated, and these values were summed to summarize population recruitment throughout task events. We first noted that A-DMS was characterized by inhibition of neurons beginning at task initiation while the P-DMS exhibited the inverse pattern (Fig.S2A-B). Furthermore, we found that both A-DMS and P-DMS exhibited widespread reductions in neuronal modulation in ‘extinction’ trials where neither port was rewarded, highlighting the relevance of this area to value-based action control (Fig.S2A-B). Taken together, our global view of population neural activity reveals: (1) A-/P-DMS populations exhibit temporally distinct modulation patterns; (2) P-DMS is predominantly activated while A-DMS is inhibited in task; (3) both regions exhibited reduced modulation when neither action was rewarded.

### A-DMS encodes previously selected action

To examine differential encoding of behavior, we focused on task-relevant variables previously found within dorsal striatum^29,30^ , starting with current trial choice, which could represent an emerging decision variable or an efference copy of the recently selected action (Fig.2I). We divided trials by ipsilateral or contralateral choice (relative to the recording site) and compared 500ms time bins against a baseline within the ITI (Fig.2J-M). We first compared neuronal modulation at each time window between A- and P-DMS, finding divergence between these populations for both ipsilateral and contralateral choice beginning roughly at task initiation in the P-DMS and beginning after choice in the A-DMS (Fig.2L, top lines). To investigate the relevant drivers of this divergence, we examined differences in population recruitment, finding significant ipsilateral versus contralateral differences after choice for both A- and P-DMS (Fig.2L,M), with a net reduction in population activity during contralateral choices in A-DMS and an increase in activity for ipsilateral choices in P-DMS. Notably the duration of this difference extended throughout the remainder of the trial for A-DMS in comparison to the transient post-choice change in P-DMS (Fig.2L,M, asterisks). To capture choice specificity at the individual unit level, we compared firing rates in ipsi-versus contralateral trials throughout the trial. The A-DMS population exhibited greater choice tuning than P-DMS throughout, with notable peaks after choice and after reward delivery (Fig.S2C,E). In contrast, P-DMS broadly exhibited less specific choice tuning except ∼1sec. prior to choice and during the ITI (Fig.S2D,F). Taken together, these data suggest temporally distinct encoding of choice along the A/P DMS axis, with persistent population representations of selected action appearing in the A-DMS immediately following action selection and a brief period of choice specificity occurring in the P-DMS both before and after choice.

### A-DMS and P-DMS encode value with differing temporal specificity

Another important behavioral variable thought to guide behavior is latent neural representations of the current value for a given action (‘action values’). Prior work has shown evidence for action value encoding in both dorsal striatum^28,31^ and its prefrontal inputs^11,32^. To estimate trial-by-trial action values, we fit a standard Q-learning model (including learning rate, inverse temperature, and bias parameters; see Methods for details) to animal choice (Fig.S3A). To test for differential representation of choice-relevant value information in the A-/P-DMS, we regressed individual neuron firing rates against action values for contralateral choice (Q_contra_) (Fig.S3B-E). Surprisingly, we found that <10% of each DMS population exhibited firing rate correlations with Q_contra_ (Fig.S3D,E). An alternative method of value representation that would be masked by regression analysis would be changes in the size of the value encoding population, such that a greater number of neurons with similar modulation would be recruited. To examine this, we separated the trials into tertiles based on Q_contra_ and compared each unit’s modulation against the ITI (Fig.3A-D). We found that both populations encoded Q_contra_ with distinct directionality and timing. Specifically, A-DMS initially exhibited a modest scaling of population inhibition with Q_contra_ that dramatically increased following choice, lasting throughout the remainder of the trial (Fig.3C). A smaller increase in A-DMS recruitment was also noted after choice and during outcome. In contrast, P-DMS representations of Q_contra_ values involved increasing unit recruitment with increasing Q_contra_ but also continued from trial initiation through outcome (Fig.3D). To further characterize the differential encoding of Q_contra_ across A- and P-DMS, we examined the relationship between the proportion of modulated neurons and their degree of modulation (defined by auROC), either before or after choice (Fig.S3F,G). For P-DMS, this analysis revealed an interesting mechanism of value encoding whereby larger Q_contra_ values were represented by larger populations of neurons with similar individual modulation levels (Fig.S3G, note the fixed location on the y-axis), a phenomenon observed both prior to and following choice selection. We performed identical analyses for neural representations of ipsilateral choice value (Q_ipsi_), again finding that A-DMS modulation strongly diverged after choice, with a sign flip such that highest Q_ipsi_ value was now associated with the greatest population suppression (Fig.S3H). In contrast, P-DMS modulation for Q_ipsi_ was in the same direction as for Q_contra_ – i.e. a greater number of units were recruited for higher Q_ipsi_ values (Fig.S3I), although here we noted smaller single unit activity modulation at the highest Q_ipsi_ values, suggesting a combination of single unit and population encoding of ipsilateral choice value (compare Fig.S3K with Fig.S3G).

**Figure 3.**
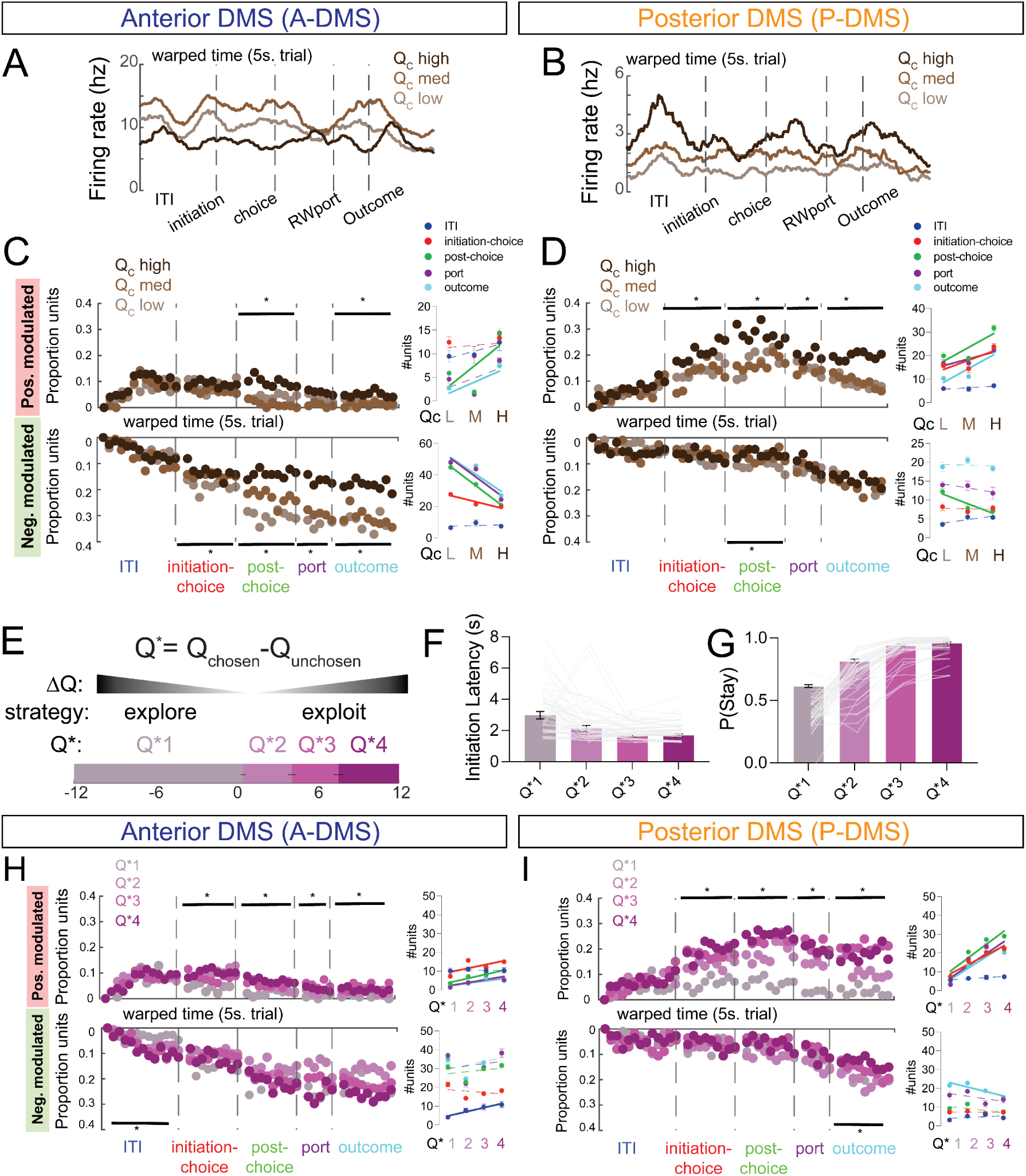
A-DMS and P-DMS encode value and strategy with differing temporal profiles. (A and B) Example striatal unit firing rates recorded in the A-DMS (A) or P-DMS (B) exhibiting modulation that scales with the action value of the contralateral choice (higher Q_contra_, darker brown). (C) Top, left: changes in the proportion of positively modulated units for different Q_contra_ values in the A-DMS. Positive modulation scales positively with value in the post-choice window and the outcome window for A-DMS. Bottom: Negative modulation inversely scales with value from the initiation-choice window through the outcome window in the A-DMS. Right panels show the average number of modulated units in each trial epoch (colored dots) as a function of Q_contra_. Solid lines represent non-zero regression values with *p<0.05, dotted lines with p>0.05. (D) Same as C, but for P-DMS. Top: positive modulation in the P-DMS scales with value from the initiation-choice window through the outcome window in the P-DMS. Bottom: Negative modulation in the P-DMS scales with value in the post-choice period only. (E) Top: Mathematical description of Q*, which is calculating by subtracting the Q-value of the unchosen port from the Q-value of the chosen port. Middle: Negative values for Q* are reflective of choices where animals chose the side with the lower value, which is a commonly used criteria for exploration. Positive values for Q* are reflective of choices where the more valued side was chosen, reflecting exploitation. Bottom: Trials were sorted by Q* and placed into quartiles (25% of trials in each bin, e.g. the bottom quartile (Q*1) accounts for all trials from ∼ Q*=-12 to +0.5). (F) Average initiation latencies within each Q* quartile show that lower values are correlated with slower times to initiate a trial, consistent with exploration. (G) Average probability of returning to a choice that was previously made within each Q* quartile, showing that higher Q* values are correlated with higher probabilities to return to a given choice, consistent with exploitation. (H) Top: changes in the proportion of positively modulated units for each Q* quartile in A-DMS. Modest increased recruitment scales with increasing Q* beginning at initiation-choice. Bottom: Modest increase in suppression with increasing Q* in the ITI window. Insets show linear regression for number of units modulated within each window according to Q*. (I) Top: changes in the proportion of positively modulated units for each Q* quartile in P-DMS. Neuronal recruitment robustly increases with increasing Q* beginning in the initiation-choice epoch. Bottom: Minimal decrease in suppression with increasing Q* restricted to the outcome window. Insets show linear regression for number of units modulated within each window according to Q*.

The differential timing and magnitude of A-DMS and P-DMS encoding of task-relevant variables suggested temporally biased functions (Fig.2I). To further test this idea, we examined the neuronal representation of current trial decision strategies, designated Q*, which is obtained by taking the difference between Q values of the chosen and unchosen options. This metric simultaneously provides information about the difficulty of the value comparison (larger absolute values representing bigger differences between the two options) and about strategy (a positive sign implying exploitation of a clear value difference and a negative sign signifying an explorational choice of the potentially lower value choice; Fig.3E). To confirm these expectations, we sorted animal behavior by Q* and examined choice patterns and execution, finding that large positive values were associated with shorter trial initiation latencies and higher probabilities of upcoming stay decisions (Fig.3F,G). We then sorted trials into quartiles according to Q* (Fig.3E) and examined population modulation in both DMS territories. Consistent with the idea that Q* represents a current-trial choice process, we found that differential population encoding of Q* in P-DMS could be first discerned between the initiation and choice, with higher Q* values correlating with a greater proportion of positively modulated units (Fig.3I). In contrast, parsing trials by this variable revealed significantly less modulation within A-DMS (Fig.3H). Together, these data further highlight the temporally distinct patterns of neuronal recruitment across the A-P DMS axis, with P-DMS strongly representing Q-values and upcoming strategy prior to choice and A-DMS providing population level information about the selected choice and its anticipated value.

### Pre-choice inhibition in the P-DMS induces a context-dependent contralateral choice bias

To investigate whether the contrasting activity patterns observed in the A-DMS and P-DMS are indicative of different functional contributions to the task, we performed inhibition of the A-DMS and P-DMS in separate mice performing the volume-comparison task. We injected AAV5-hSyn-NpHR3.0-EYFP and implanted 200um fiber optic cannulas into the A-DMS or P-DMS bilaterally (Fig.4A). We first examined the effects of inhibition prior to choice, delivering continuous light from the beginning of a trial until the choice was registered (designated as ‘pre-choice inhibition’) (Fig.4A). As our logistic regression analysis revealed an extended integration window of prior reward history (Fig.S1B), we elected to perform inhibition on every trial in a randomly selected 50% of blocks. Further, given the strong contralateral control of behavior within striatum^33–35^, we initially performed unilateral manipulations, defining the optogenetic target site in relation to the more valued side (‘optimal’; Fig.4B,E). We found that there was no effect of unilateral pre-choice perturbation in the A-DMS on choice, accuracy, probability of staying, task latencies, or ‘extinction’ behavior, either in the 12v0 (Fig.S4B-H) or 12v4 blocks (Fig.4C,D). In contrast, we found that pre-choice inhibition in the P-DMS aided performance when the high-value side was contralateral to the manipulation and impaired performance when the manipulation was ipsilateral to the high-value side (Fig.4F,G), consistent with a bias towards choosing the port contralateral to the manipulation site (Fig.S4N). Notably, we found that this effect was apparent in 12v4 blocks but not 12v0 blocks (Fig.S4I,J). We did not observe any impact on motor execution or ‘extinction’ performance (Fig.S4K-O). Furthermore, these effects were not seen in GFP control mice with A-DMS or P-DMS manipulations (Fig.S5). To rule out that the observed contralateral bias arose from a simple motor or ‘body orienting’ effect, we performed unilateral inhibition of the P-DMS as mice explored an open field (Fig.S6). We compared the impact of optogenetic stimulation on velocity and body angle at laser onset and offset, noting no significant differences aside from a small velocity increase following bilateral inhibition (Fig.S6C-J).

**Figure 4.**
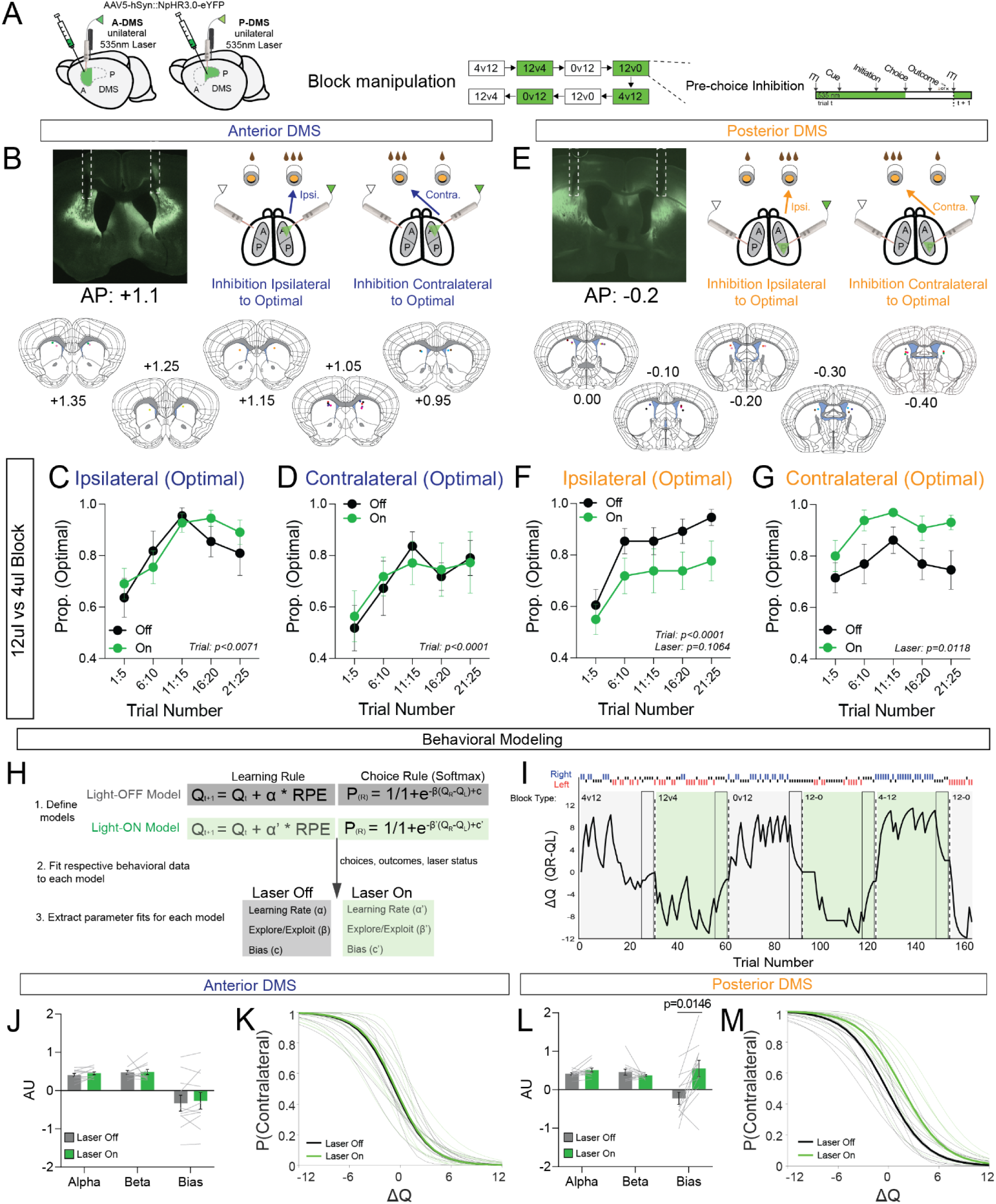
Unilateral pre-choice inhibition induces a contralateral bias in the P-DMS, but has no effect in the A-DMS. (A) Experiment schematic. Unilateral activation of NpHR3.0 in the A- or P-DMS in 50% of blocks. Every trial in a ‘light on’ block gets continuous 532nm light delivery from ITI to choice selection. (B) Representative histological section showing virus and fiber tract in the A-DMS (top left). Light delivery schematic shows the relationship between the optogenetic manipulation and the optimal choice side (top right). Fiber tract tip location mapped for animals mapped throughout the A-DMS (bottom). (C) Proportion of optimal choices in 12v4 blocks when the laser is ipsilateral to the optimal choice and is off (black) or on (green) (*n*=11; RM-two-way ANOVA, trial: F_2,18_ = 7.000, **P = 0.0071, Laser: F_1,10_ = 0.8164, P = 0.3875, interaction: F_2,21_ = 0.8864, P = 0.4327). (D) Same as C, but for 12v4 blocks when the laser is contralateral to the optimal choice (*n*=11; RM-two-way ANOVA, trial: F_2,24_ = 12.09, ***P = 0.0001, laser: F_1,10_ = 0.009207, P = 0.9255, interaction: F_2,20_ = 0.3031, P = 0.7423). (E) Same as B, but for P-DMS. (F) Same as C, but for P-DMS (*n*=12; RM-two-way ANOVA, trial: F_3,29_ = 17.23, ****P < 0.0001, laser: F_1,11_ = 3.093, P = 0.1064, interaction: F_2,21_ = 0.3031, P = 0.8326) (G) Same as D, but for P-DMS (*n*=12; RM-two-way ANOVA, trial: F_2,20_ = 2.286, P = 0.1317, laser: F_1,11_ = 9.066, *P = 0.0118, interaction: F_2,18_ = 0.5725, P = 0.5412) (H) Q-Learning model with separate parameter fits for ‘laser-on’ and ‘laser-off’ trials. (I) Representative plot of individual animals ΔQ values during an optogenetic session. Block volume comparisons are in upper left (green shading for ‘laser-on’ blocks) and animals’ right and left choices are in blue and red. (J) Average model parameters for A-DMS mice are not different for laser-off and laser-on blocks (*n*=11; Wilcoxon matched pairs rank sum test with Holm-Šídák correction for multiple comparisons, Alpha: P = 0.744236, Beta: P = 0.769150, Bias: P = 0.769150). (K) Individual (thin lines) and average (thick lines) softmax choice fits for A-DMS animals in laser-on (green) and laser-off (black) conditions. (L) Average model parameters for P-DMS mice show a higher bias to contralateral choices for laser-on trials compared to laser-off trials (*n*=12; Wilcoxon matched pairs rank sum test with Holm-Šídák correction for multiple comparisons, Alpha: P = 0.242046, Beta: P = 0.469727, Bias: *P = 0.014577). (M) As in (K) for P-DMS animals.

To better understand how pre-choice manipulations altered value-based choice behavior, we used parallel Q-learning reinforcement learning models whereby latent variables were fit independently for control and optogenetic manipulation trials, across both block types (Fig.4H,I). We then compared parameters describing the value updating (alpha), choice function (beta) and bias terms of the RL model. While we did not observe any differences in the fits for A-DMS (Fig.4J,K), in the P-DMS we noted a significant change in the bias term (Fig. 4L), indicative of a bias to contralateral choice in optogenetic trials. Plotting the choice probability (Fig.4M) as a function of ΔQ (the difference between Q_ipsi_ and Q_contra_) for the P-DMS showed that the impacts of optogenetic manipulation were strongest around smaller absolute ΔQ values, with minimal effects when either the ipsilateral or the contralateral choice were clearly more valuable. These data suggest that pre-choice P-DMS activity has a context-dependent modulatory function on choice. To see whether this could account for the differing impacts of manipulation depending on block type, we separated trials from each block into three bins: trials where the choice contralateral to manipulation is much higher value than ipsilateral (ΔQ = -12 to -4), trials where the choice ipsilateral to the manipulation is much higher value than contralateral (ΔQ = 4 to 12), and trials where the value of the two choices are closer in value and therefore more difficult to distinguish (ΔQ = -4 to 4). We found that the effect of inhibition was strongest in ambiguous value environments (i.e. ΔQ = -4 to 4) (Fig.S4R,S). This, together with the larger proportion of challenging value choices found in 12v4 blocks (Fig.S4R,S, bottom), explains the weaker optogenetic effects of P-DMS manipulation in 12v0 contexts. Consistent with the unaffected choice patterns, we did not observe any optogenetic-dependent differences when our behavioral models were applied to A-DMS (Fig.4J,K; Fig.S4P,Q).

Given prior descriptions of the lateralized impact of striatal function^33–35^, we examined whether bilateral pre-choice inhibition in the A-DMS or P-DMS had distinct effects from ipsilateral manipulation using a similar block-wise protocol. Consistent with our data, we did not see any effects of bilateral pre-choice inhibition on accuracy, extinction performance, or task latencies in the A-DMS (Fig.S7B-D, H-J). In contrast, while bilateral pre-choice inhibition in the P-DMS did not change choice accuracy or extinction behavior (Fig.S7E-G), we observed an increase in latency to initiate and register choice, leading to a reduction in the overall number of completed trials per fixed session length (Fig.S7K-M). These effects on latency were not observed in GFP controls (Fig.S8). Together, our data suggest that pre-choice manipulation of the P-DMS can bias current trial choice contralaterally in a value-dependent manner. In contrast, bilateral manipulations do not impact choice but rather delay execution of motor output. Furthermore, pre-choice perturbations of A-DMS do not impact choice or motor execution.

### Unilateral post-choice inhibition in the A-DMS impairs contralateral reward processing

Next, we looked at how post-choice manipulation of A-DMS and the P-DMS activity affected task performance. The dynamic population reorganization following choice in the A-DMS and the persistent population differences accumulating before choice in the P-DMS suggested that both areas could contribute to the evaluation of selected actions. To test this, we activated halorhodopsin beginning immediately after choice selection and throughout the outcome delivery on every trial in 50% of blocks (Fig.5A, designated ‘post-choice’). Surprisingly, in the A-DMS we found that inhibition affected choice when the high-value port was contralateral to the manipulation (Fig. 5B,C). Consistent with earlier pre-choice manipulations, these effects were noted only in the 12v4 blocks (Fig.S9B-D). In contrast, there was no major effect of inhibition in the P-DMS in either 12v0 or 12v4 blocks (Fig.5D,E; Fig. S9I-K) and no effects on latency or overall choice bias were observed in either condition (Fig.S9E-H, L-O). Finally, none of these effects were observed in GFP control mice (Fig.S10).

**Figure 5.**
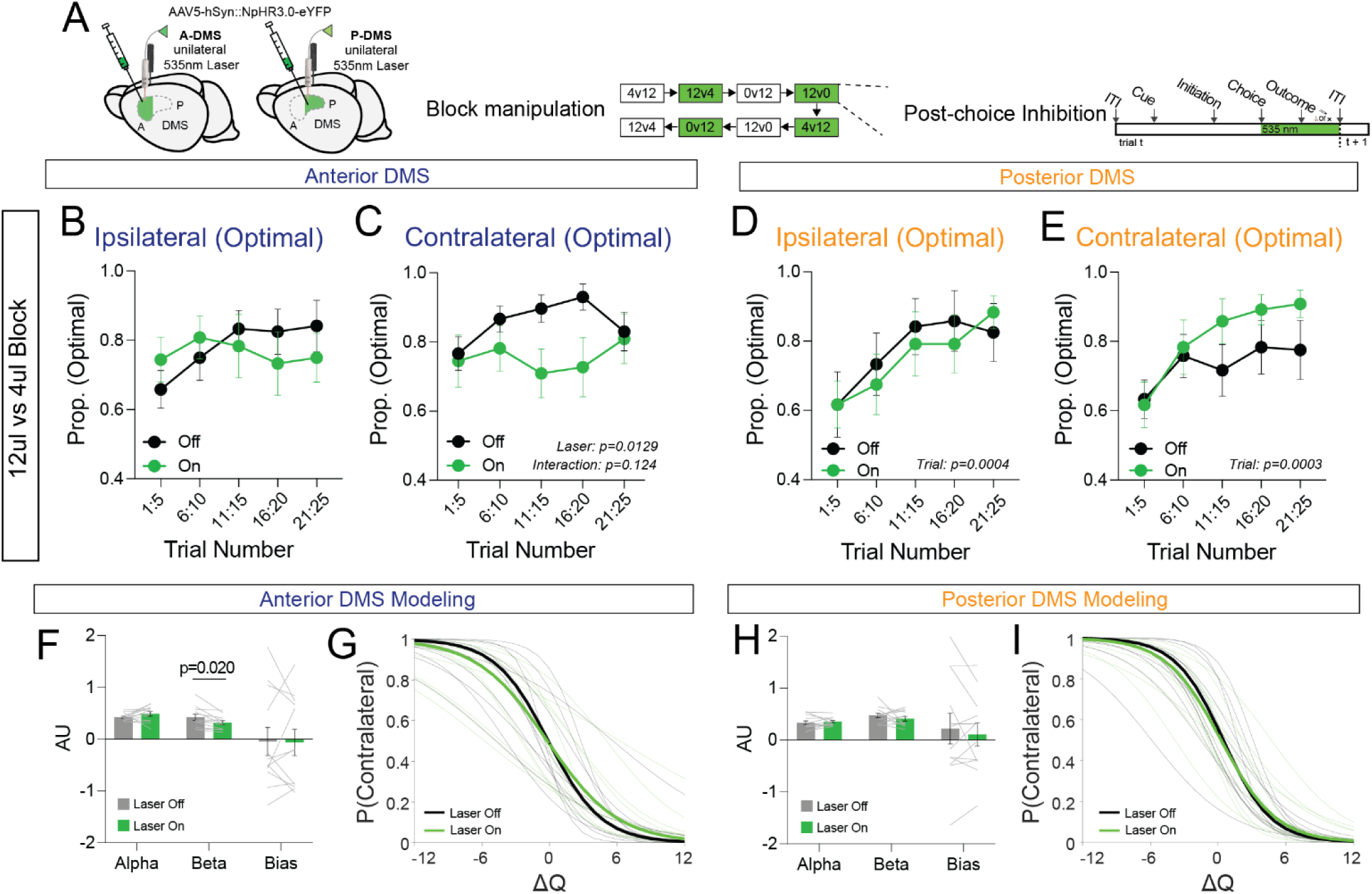
Unilateral post-choice inhibition in the A-DMS impairs subsequent reinforcement, but has no effect in the P-DMS. (A) Experiment schematic. Unilateral activation of NpHR3.0 in the A- or P-DMS in 50% of blocks. Every trial in a ‘light on’ block gets continuous 532nm light delivery from choice to ITI start. (B) Proportion of optimal choices made by A-DMS animals in 12v4 blocks when the laser is ipsilateral to the optimal choice and off (black) or on (green) (*n*=11; RM-two-way ANOVA, trial: F_2,25_ = 1.435, P = 0.2576, laser: F_1,11_ = 0.1406, P = 0.7148, interaction: F_2,25_ = 0.9762, P = 0.4003). (C) Same as B, but for 12v4 blocks when the laser is contralateral to the optimal choice (n=11; RM-two-way ANOVA, trial: F_2,19_ = 0.4595, P = 0.6528, laser: F_1,10_ = 9.126, *P = 0.0129, interaction: F_2,23_ = 2.226, P = 0.1241). (D) Proportion of optimal choices made by P-DMS animals in 12v4 blocks when the laser is ipsilateral to the optimal choice and off (black) or on (green) (*n*=12; RM-two-way ANOVA, trial: F_3,32_ = 8.116, ***P = 0.0004, laser: F_1,11_ = 0.0675, P = 0.8099, interaction: F_2,22_ = 0.5557, P = 0.5840). (E) Same as D, but for 12v4 blocks when the laser is contralateral to the optimal choice (n=12; RM-two-way ANOVA, trial: F_3,37_ = 7.732, ***P = 0.0003, laser: F_1,11_ = 2.273, P = 0.1598, interaction: F_3,35_ = 0.8766, P = 0.4671). (F) Average model parameters for A-DMS mice show a reduced beta for laser-on compared to laser-off blocks (*n*=11; Wilcoxon matched pairs rank sum test with Holm-Šídák correction for multiple comparisons, Alpha: P = 0.321468, Beta: *P = 0.020368, Bias: P = 0.850098) (G) Individual (thin lines) and average (thick lines) softmax choice fits for A-DMS animals in laser-on (green) and laser-off (black) conditions. (H) Average model parameters for P-DMS mice are not different for laser-off and laser-on blocks (*n*=12; Wilcoxon matched pairs rank sum test with Holm-Šídák correction for multiple comparisons, Alpha: P = 0.711661, Beta: P = 0.718810, Bias: P = 0.850098). (I) As in (G) for P-DMS animals.

These results suggest a unique role for A-DMS in processing reward outcomes in a side-dependent manner to influence future choice. We again used a modified Q-learning RL model to estimate what drove the observed inefficient choice behavior (Fig.5F-I). In contrast to pre-choice perturbation of P-DMS, we found that A-DMS animals had a lower beta parameter for laser-on trials, indicating that animals were less likely to use reward history to guide choice when the laser was on. (Fig. 5F,G). There were no alterations to model parameters seen in post-choice P-DMS manipulation (Fig. 5H,I). Finally, we examined the effects of bilateral post-choice inhibition.

We found that bilateral post-choice inhibition in the A-DMS paradoxically improved performance, an effect that was again seen in 12v4 but not 12v0 blocks (Fig.S11). In contrast, bilateral post-choice inhibition in the P-DMS had no effect on choice, accuracy, or latency (Fig.S11). No effects were observed in GFP control mice receiving bilateral post-choice inhibition in either striatal region (Fig. S12). These data suggest that optogenetic perturbations immediately following choice in the A-DMS, but not the P-DMS, interfere with neural signals used to evaluate and select future choices. Together, our perturbations resolve distinct regional DMS functional contributions for the selection and evaluation of value-based choice.

### A-DMS and P-DMS share common outputs in the ventral medial SNr, which exhibits transformed value representations present in both striatal areas

Given prior work showing an integration of cortical areas within the A-/P-DMS, we asked whether there was similar convergence of these striatal regions in the main basal ganglia output nucleus, the substantia nigra, pars reticulata (SNr) . To examine this, we first performed anterograde trans-synaptic tracing^35,36^ of SPNs projecting to SNr by injecting AAV1-hSyn-Cre (N=8) or AAV1-Ef1α-FLP (N=8) into the A-DMS and P-DMS, respectively, together with injection of AAV5-flex-mCherry and AAV5-fDIO-EYFP viruses throughout the SNr (Fig.6A-C). With this approach, the postsynaptic SNr targets of direct pathway A-DMS neurons will be labeled with mCherry while P-DMS SNr targets will be labeled with EYFP (Fig.6C,D). We found that A-DMS projections were biased to the ventromedial SNr (vmSNR) throughout its A-P extent. In contrast, P-DMS projections are more diffusely distributed across the mediolateral and dorsal-ventral SNr axes, with distinct patterns along the A-P extent (Fig. 6D,E)^37^. Thus, while A-DMS and P-DMS largely target distinct parts of the SNR, the vmSNr represents a potential site of convergence.

**Figure 6.**
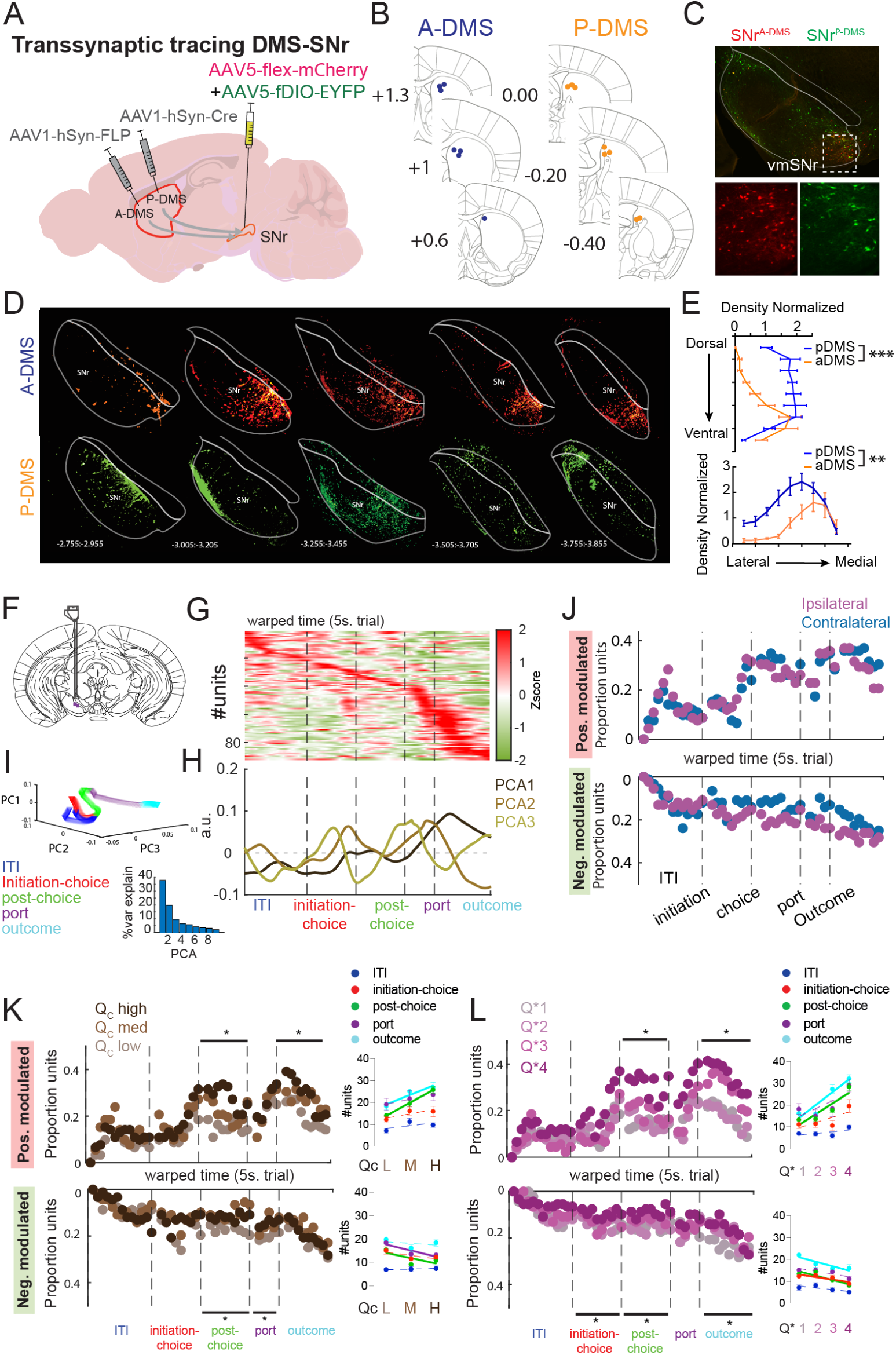
Anatomical and functional relationships between SNr and DMS subregions. (A) Anterogradely-transported AAV1-hSyn-FLP and AAV1-hSyn-Cre were injected into the A-DMS and P-DMS, respectively, while Cre- and FLP-dependent reporter viruses (mCherry and EYFP, respectively) were injected throughout the SNr (N=8), thereby labeling SNr neurons downstream of A- and P-DMS. (B) Anatomical localization of AAV1 viruses in the A-DMS (left) and P-DMS (right). (C) Top: representative image of SNr showing distribution of red (A-DMS targeted) and green (P-DMS targeted) SNr cell bodies. White box marks enlarged vmSNr area below. Bottom: Split channels for A-DMS targeted (left) and P-DMS targeted (right) SNr neurons. (D) Mapping of SNr neurons that receive projections from the A-DMS (top, red) or P-DMS (bottom, green) across the anterior-posterior SNr axis. (E) SNr neurons that receive projections from the A-DMS are consistently localized in ventral-medial parts of SNr. In contrast, SNr neurons that receive projections from the P-DMS are localized more diffusely in the dorsoventral and mediolateral axes, depending on A-P location. (F) Electrode placement in the vmSNr for recording isolating extracellular units (N=2). (G) Averaged activity of vmSNr units recorded in the volume block task (time warped) aligned to trial start. Units are sorted by peak activity. (H and I) The first three principal components of averaged vmSNr activity highlights peaks in signal variability following choice selection. (J) Top: Positive modulation of vmSNr neurons across the task for ipsilateral and contralateral choices. vmSNr neurons were recruited during the ITI and following choice, however, no differences in recruitment between ipsilateral and contralateral choices were noted. Bottom: Negative modulation of vmSNr neurons for ipsilateral and contralateral choices. SNr neurons were negatively modulated throughout the trial, however, no differences in recruitment between ipsilateral and contralateral choices were noted. (K) Top: changes in the proportion of positively modulated units at different Q_contra_ values in vmSNr. Positive modulation increases with increasing Q_contra_ value in the post-choice and outcome windows for vmSNr. Bottom: Negative modulation decreases with increasing Q_contra_ value in the post-choice and port-return windows in vmSNr. Insets show linear regression for number of units modulated within each window according to Q_contra_ value. (L) Top: changes in the proportion of positively modulated units for each Q* quartile in vmSNr. Positive modulation increases with more positive Q* in the post-choice and the outcome windows. Bottom: Negative modulation weakly scales with Q* in the initiation-choice, post-choice, and outcome windows. Insets show linear regression for number of units modulated within each window according to Q*.

To examine whether this SNr domain similarly encoded aspects of value-based choice, we performed extracellular recordings from the vmSNR, identifying a total of 90 neurons from mice (N=2) engaged in the volume-comparison task (Fig.6F). We examined average single unit activity via PCA, finding that the weights of the first three components (explaining ∼65% of the total variance) exhibit peaks just prior to choice and as the mice entered the port and experienced outcomes (Fig.6G-I). The predominant pattern of activity was that of increased unit recruitment following choice and lasting until the next trial, with a small, consistent increase in unit suppression (Fig.S13A). The vmSNr exhibited reduced positive neuronal modulation but no changes in unit suppression during ‘extinction’ trials where neither side was rewarded (Fig.S13A). This was unexpected given the SNr’s well-documented role in movement, suggesting a role for the vmSNR in goal-directed action and value-seeking behavior.

We then investigated the neuronal representation of choice in the vmSNr, finding an increased recruitment of neurons beginning just prior to choice and lasting through the remainder of the trial (Fig 6J). While echoing the pattern of A-DMS, vmSNr did not distinguish between ipsilateral and contralateral choices at the population level. We did however note elevated choice tuning at the single unit level following choice selection (Fig.S13B,C). To examine whether the vmSNr contained value-related neural signals, we first focused on Q-value encoding. While the post-choice timing of population neuronal modulation was more aligned with the A-DMS, the encoding of Q values in vmSNr exhibited greater similarity to P-DMS along multiple dimensions – (1) at the population level, both increasing Q_contra_ and Q_ipsi_ were encoded by increasing numbers of vmSNr units (Fig.6K, Fig.S13E); (2) at the single unit level, neurons recruited for Q_contra_ exhibited similar firing rate modulation while those recruited for higher Q_ipsi_ exhibited smaller individual modulation scores (Fig.S13F). Finally, we examined whether vmSNr activity encoded current trial strategy, using Q* values. We found that vmSNr exhibited the largest single unit recruitment increases for exploitational choices, a pattern robustly observed in P-DMS (Fig. 6L). Taken together, these data suggest that the vmSNr, a region targeted by both the A- and P-DMS, exhibit unique neural representations for value-based choice that combine aspects of A- and P-DMS region encoding.

## DISCUSSION

Prior lesion studies have suggested divergent functions along the A/P axis^15–17^ while analysis of prefrontal cortical circuits defined by A/P striatal target show a clear segregation of behaviorally relevant neural signals^11^. Utilizing a systematic recording and perturbation approach, we sought to identify a unifying functional architecture of the dorsomedial A/P axis and its downstream output connectivity within SNr. Foremost, we found widely divergent population neural dynamics according to A/P site within DMS, with inhibition and recruitment predominating in

A-DMS and P-DMS, respectively (Fig.2). In addition, we reliably noted significant reductions in neuronal modulation in ‘extinction’ trials when neither port was rewarded. Regarding key neural variables for value-based choice, we found that A-DMS most strongly represented the choice and its anticipated value immediately following choice selection, while the P-DMS encoded action values and strategy preceding choice selection. Consistent with these temporal differences, we found that pre-choice unilateral optogenetic perturbation of P-DMS contralaterally biased upcoming choice while post-choice perturbation of A-DMS reduced reinforcement of choice when the optimal side was contralateral to our manipulation. Interestingly, both manipulations only impacted behavior in blocks with more challenging value comparisons, suggesting context-specific involvement. Finally, we showed that despite largely projecting to distinct SNr regions, the A- and P-DMS were integrated in the vmSNr, a site whose neurons exhibited robust encoding of key value-based behavioral parameters. Taken together, our data suggest a temporally distributed model for the selection and evaluation of choices along the A-P striatal axis and its downstream targets.

### Divergent Neuronal Representations of Value-Based Choice Across the Anterior-Posterior DMS Axis

While population level analysis of neural activity revealed numerous differences along the A/P striatal axis, the most striking difference was the consistent directionality – most P-DMS modulation involved recruitment while the majority of A-DMS modulation exhibited suppression of neuron firing rates. The local circuit mechanisms supporting this divergence remain unclear. One interesting possibility relates to greater use of feed-forward inhibitory architectures within the A-DMS, which could mediate the type of immediate inhibition occurring at choice. Studies of prelimbic (PL) neurons targeting A- and P-DMS demonstrated that these projections specifically recruited extensive feed-forward inhibition in A-DMS, while recordings during a cost-benefit task highlighted the importance of fast-spiking interneuron-mediated feed-forward inhibition in decision processes^11,38^. The relevance of this circuit design awaits a better understanding of the local circuit interneuron elements that generate feed-forward inhibition in A-DMS. Another global trend in our recording data was that ‘extinction’ trials, where neither port was rewarded, exhibited significant reductions in neuronal modulation, both in the DMS and the vmSNr. These data are particularly striking given that the animals continued to perform the same action patterns, highlighting the circumscribed importance of DMS circuits within value-based choice environments.

Neural representations of choice can represent either an emerging decision process or a retrospective accounting of the recently selected choice. Interestingly, we found a divergence in encoding of choice along the A/P DMS axis. In A-DMS, we observed a robust network reorganization immediately following choice selection, with a net effect of reduced population activity for contralateral but not ipsilateral choices throughout the remainder of the trial. In contrast, P-DMS exhibited a smaller and temporally restricted increase in unit activity following ipsilateral choice. A comparison of individual neuron choice tuning revealed further region-specific differences – A-DMS exhibited broader choice-specific tuning with peaks following choice and outcome, while P-DMS had discrete tuning epochs immediately prior to choice (Fig.S2E,F). These data suggest a potential role for the A-DMS in monitoring of selected actions by providing an ‘efference trace’ signal to downstream circuits involved in linking outcomes with future strategies. The origins of these A-DMS signals remains unknown but models of oculomotor learning have posited that pre-motor areas responsible for the action also use collateral striatal connections to gate plasticity associated with reward signals^39^. Our prior work has found that PL projections targeting the A-DMS were uniquely enriched in encoding of recently selected action, although the observed phasic responses cannot fully account for the sustained population response seen in our A-DMS recordings.

Value-based decisions require the real-time tracking of value and its transformation into action strategies. Choice values and strategy were differentially represented along the A/P DMS axis, with interesting differences in the timing and manner of modulation. For example, contralateral Q-value encoding began at trial initiation in P-DMS but was most pronounced only following choice selection in the A-DMS. Furthermore, increasing Q_contra_ was represented in P-DMS via increases in the number of recruited units as opposed to scaling of individual unit firing rates. This contrasts with Q_ipsi_ encoding within P-DMS, where higher action values recruited more units that nevertheless had weaker overall rate modulations (compare Fig.S3K with Fig.S3G). Together, these data suggest a new population-based framework for our understanding of value representation within striatal populations. To assess strategy encoding, we examined Q*, a behavioral parameter that describes Q_contra/ipsi_ contrasts and their relationship to next-trial choice, defined as either exploitation (selecting the higher Q option) or exploration (picking the lower Q option). Consistent with P-DMS encoding information relevant for the upcoming choice, we found that exploitation of obvious choice value differences was associated with a large P-DMS recruitment starting at trial initiation. The relationship between Q* and A-DMS activity was in contrast weak. Taken together our neural recordings reveal highly divergent patterns of modulation according to A/P DMS location, with neural signals for choice selection in P-DMS and choice monitoring in A-DMS. In our attempt to address striatal regional function, we did not consider SPN subtype in either recordings or neural perturbations. While 1-photon calcium imaging has shown similar representations of choice tasks across dorsal striatal output pathways^29^, other studies have clearly highlighted the importance of considering this in future work^19,30,35,40^.

### Temporally Distinct Control of Goal-Directed Behavior across the Anterior-Posterior DMS Axis

Our neural recordings made clear functional predictions that we tested with non-cell type specific optogenetic inhibition, probing both the timing (before or after choice) and laterality (unilateral versus bilateral) of effects. We found that unilateral P-DMS inhibition increased the selection of contralateral choice in 12v4ul but not 12v0 blocks. Further analysis suggests this context-dependence relied on the fact that 12v4 blocks have a greater proportion of trials with small value contrasts between choices (see ΔQ = -4 to 4 in Fig.S4S), the value space where light had its strongest choice effects. Consistent with these data, we found that this change in performance was captured by an increase in the bias term of our model, creating a lateral shift in the psychometric curve for ΔQ’s impact on choice when the laser was on. This shift has the largest effect in the middle of the curve and little effect on the extreme ΔQ values. This dependence of striatal relevance on task demands is consistent with prior work using multiple virtual T-maze task variants^41^, and highlights the dynamic flexibility of the DMS to motor control. It is important to note that we did not observe differences in neural activity in P-DMS according to reward block, suggesting that this diversion from DMS control is not correlated with reduced neuronal recruitment. Our data are also generally consistent with prior studies showing optogenetic excitation of dSPNs mimicking a transient increase in ‘action value’ of contralateral targets^42^. While our manipulation would impact both SPN subtypes, it is possible that iSPN-mediated inhibition leads to a sustained suppression of SNr via GPe release that cumulatively overrides any impacts of dSPN dis-inhibition of SNr. Whether simultaneous inhibition would lead to this net thalamic dis-inhibition will require further testing. Finally, it is important to highlight the absence of any impacts on choice when both striatal hemispheres were simultaneously inhibited. It was however accompanied by a general slowing of motor performance, suggesting that bilateral P-DMS activity supports the invigoration of choice performance. These data together with others^11,15,16^ confirm the function of P-DMS for value-dependent biasing and execution of choice.

Consistent with the larger population changes observed following choice in the A-DMS, our post-choice manipulations were exclusively effective in this region. Despite the differences in timing, there were several similarities to our P-DMS results, both regarding the context-dependence of optogenetic effects and the contralateral nature of the observed phenotype. The select phenotype in 12v4 blocks (as seen in P-DMS) further reinforces the idea that these value-related circuits can be circumvented dependent on task demands. The reduction in contralateral accuracy when A-DMS was unilaterally inhibited following choice is suggestive of a role in choice monitoring and evaluation. We found that this change in ability to use previous trial information to determine future choice was captured by a reduction in the beta term of our behavioral model when the laser was on. It’s possible we might have found a larger change in beta if not for the ipsilateral trials, which had no phenotype for inhibition, being governed by this same parameter. Given that the reward outcome is broadly experienced throughout the mouse’s brain, this lateralization may reflect one neural circuit mechanism for restricted credit assignment^43^. Given the unique organization of afferent excitatory inputs along the A-P axis, wherein A- and P-DMS projections are closely intermingled^11^, it is tempting to hypothesize that neural activity that shapes choice in P-DMS can be directly relayed to A-DMS-projecting populations to represent the selected action and its expected value. Whether this signal originates from cortical or thalamic projections and how it might be combined with outcome to influence subsequent choice will require further study. The use of a lateralized accounting mechanism for value-based choice may also help explain the puzzling contralateral neglect arising from unilateral striatal lesions or glutamate antagonism^44,45^. Another interesting parallel with P-DMS manipulations is the difference in behavioral effects between unilateral and bilateral manipulations. Bilateral A-DMS inhibition instead produced improved choice accuracy seen in 12v4 but not 12v0 blocks. While the exact mechanisms of this paradoxical improvement are unclear, these data are broadly consistent with SPN subtype-specific optogenetic inhibition studies in A-DMS, wherein dSPN suppression facilitates reversal learning^21,46^.

### vmSNr Integrates Value-Based Information from the Anterior-Posterior DMS Axis

Our anterograde anatomical tracing studies of SNr connectivity from A-/P-DMS are consistent with prior work showing spatially resolved targeting dependent on region of striatal origin^37^. However, our simultaneous injections have provided further insights – while aDMS projections reliably targeted the vmSNr, we found that neurons downstream of P-DMS were more broadly distributed throughout SNr with different targeting patterns along it’s rostral-caudal extent. Furthermore, the vmSNr was the only site where SNr neurons downstream of both A-/P-DMS could be found, suggesting a potential site of integration between these striatal areas. While further electrophysiology is needed to quantify the degree of direct synaptic convergence on individual SNr neurons, their wide-spread local axon collateralization may be sufficient to mediate integrative processing from these two striatal areas^47^.

Previous accounts of SNr function have been intimately linked to movement, including control of saccadic gaze^48,49^, regulation of reach kinematics^50^, and positional turning^51^. While these studies arise from multiple sub-regions of SNr, optogenetic manipulations of medial SNr have been shown to increase gait asymmetries^51,52^. Surprisingly, our recordings in vmSNr did not provide strong evidence of movement related signals. This claim is supported by multiple pieces of evidence: (1) neural representations of current trial choice are not seen during execution but only after choice is selected; (2) ‘extinction’ trials where mice performed similar motor sequences as in rewarded trials exhibited blunted positive recruitment of vmSNr; (3) optogenetic inhibition of the A/P-DMS regions upstream of vmSNr did not produce detectable motor effects. Further manipulations of SNr populations specifically downstream of DMS will help to resolve any specific contributions of vmSNr to the orienting responses previously observed. In contrast to the weak encoding of movement, we observed robust neural representations of choice value and strategy in the vmSNr, consistent with their anatomical connectivity to DMS. Surprisingly, scaling of neural activity with Q_contra_, Q_ipsi_, and Q* were similar to the encoding of these variables in the P-DMS while the temporal patterns were more similar to A-DMS, particularly in their lack of differential modulation pre-choice. One possibility is that pre-choice related information originating in the P-DMS, which we show both biases choice and shapes motor performance, may be routed through one of this striatal region’s numerous other SNr targets. If so, future work should consider whether non-motor related neural activity originating in P-DMS that targets vmSNr may contribute to the evaluation of choices or policy shifts. Overall, these data contribute to a small but growing body of work on value-encoding within the SNr and highlight value-related post-choice signals in mouse vmSNr. SNr GABAergic unit recordings during a timed pressing task have revealed neural representations for confidence of the recently completed action and reward prospects^24^, signals akin to the evaluative ones we find in our recordings. In sensory-guided choice, the long-term value of objects associated with varying reward volumes is stably represented in the caudo-dorsolateral SNr, a region downstream of the caudate tail in non-human primates^26^. Furthermore, similar value and certainty signals were also found in vlPFC, another node in this basal ganglia circuit, and activity in both regions paralleled free gaze bias^25^.Together, these data highlight the growing appreciation of value representations distributed throughout the SNr.

Overall, our data suggest there is temporally segregated processing of value-based choice across the anterior-posterior axis of the DMS, with posterior regions guiding present choice and anterior regions evaluating and updating future actions. These signals are potentially integrated in the vmSNR to exert control over regions that gate movement and decision-making.

## METHODS

### Animals

Animal experiment procedures were approved by the University of Pennsylvania Institutional Animal Care and Use Committee, and all experiments were conducted in accordance with the National Institutes of Health Guidelines for the Use of Animals. Animals were acquired from Jackson Laboratory, (c57/bl6, strain 000664) were grouped with littermates on a 12:12 light-dark cycle and provided *ad libitum* food and water. Given the potential for impacts of the estrous cycle on goal-directed behavior, all experiments were conducted on naive male mice.

### Behavioral equipment

Behavioral experiments were performed in a custom built 3-port operant chamber (Sanworks LLC, NY). Each port has a white LED light, an infrared beam emitter, an infrared beam detector, and an outlet for liquid reward delivery. The center port was designated as a reward delivery outlet using a pinch valve (225P011-21, NResearch, NJ). All behavioral chambers were enclosed in sound-attenuating boxes (PSIB27, Pyle, NY). Behavior protocols were controlled by Bpod software (https://github.com/sanworks/Bpod) in MATLAB (MathWorks). All port entries and events were recorded by the Bpod State Machine during behavioral sessions.

### Stereotaxic surgery

Intracranial surgery was conducted on a stereotaxic surgery frame (Kopf Instrument, Model 1900) under isoflurane anesthesia (1.5–2% + oxygen 1 L/min). Animal body temperature was maintained at 30 °C during surgery using a feedback thermocontroller (Harvard apparatus, #50722 F). Skin was cleaned with Nair hair remover followed by application of betadine to disinfect the area. Prior to surgery, 2 mg/kg bupivacaine was administered subcutaneously, and the mouse was given a single dose of meloxicam (5 mg/kg). Skin was carefully opened along the anterior-posterior midline, bregma was set to zero based on skull balance. A craniotomy was performed with a drill above the target site. Virus or Tracer was loaded into mineral oil (Sigma-Aldrich, M3516)-filled glass pipette (WPI, TW100F-3) and delivered at rate 30 nl/ min using a micro-infusion pump (Harvard Apparatus, #70-3007). At least 5 minutes after infusion, the pipette was slowly withdrawn (1 mm/min) from the brain, and the skin was sutured. Animals were monitored up to 1 h following regaining of consciousness, then transferred to the home cage and monitored after 24 h, 48 h and 72 h. Injection coordinates, A-DMS: AP +1.15 mm, ML +1.35 mm, DV −2.8 mm; P-DMS: AP −0.25 mm, ML +1.9 mm, DV −2.7 mm.

### Behavioral training

Mice were calorie-restricted to 85-90% of their *ad libitum* weight in the week prior to training. This weight level was maintained throughout the operant training and all behavioral experiments to promote operant responding. Initial training involved illuminating the center port of the behavioral chamber and offering a liquid reward whenever the animal poked into it. After reaching a criteria of poking 10 times a minute, animals progressed to light-guided sessions where the illuminated center port served as a trial initiation. Following initiation, one of the two side ports was randomly illuminated. If animals poked into the illuminated side-port, they received a liquid reward in the center port. Once animals could perform the light-guided sessions with >80% accuracy, they were moved onto training in a two-alternative forced choice task. This training task has been published in our lab prior^11^. In brief, the task involves (1) an illuminated center port that the animal pokes into to initiate a trial. (2) Both side-ports are illuminated and the animal makes a choice by poking into one of them. (3) The animal pokes into the center port to receive the outcome of their choice (reward or no reward). Only one side is rewarded on any given trial and the rewarded side switches after the animal has chosen the rewarded side on 8/10 trials in a 10-trial moving window. There is a 3 second ITI between trials. When animals reached the criteria of 75% accuracy and 200 trials per hour, they were moved onto the volume comparison task.

### Two-alternative volume comparison task

The structure of a trial in the volume comparison task is as follows: (1) The center port is illuminated at the start of a trial. Animals can initiate the trial by nose-poking into the illuminated port. (2) The two lateral choice ports are illuminated. Animals make a choice by poking into either of the two ports. If they do not register a choice within 10 seconds, the trial ends and the ITI begins. (3) After the animal makes a choice, the side port lights are turned off and the center port is illuminated. The animal must poke into the center port to receive their reward. There is a variable 50-150ms delay after the animals poke into the center port before the pinch valve opens to dispense reward. (4) The reward consumption phase continues until the animals exit the port. (5) There is a 1 second ITI between trials where all lights are extinguished. On a given trial, there is an optimal choice port that gives 12ul of reward 70% of the time. The other 30% of trials are considered ‘noise’ trials where animals do not receive any reward. The other port will be suboptimal and will deliver either 4ul of reward (in 12/4 blocks) or 0ul of reward (in 12/0 blocks). In 12/4 blocks, the suboptimal port will give 4ul of reward 70% of the time and the other 30% of trials are considered ‘noise’ trials. Each block of 12/0 or 12/4 will last 25 trials. There are 5 depreciation trials between each block where both sides are unrewarded to prevent animals from developing a bias toward any port. At the start of the first block, the optimal side is randomly assigned. For every subsequent block, the optimal side will alternate with the suboptimal side. Animals were trained in this task daily for hour-long sessions until they chose the optimal choice 75% of the time and completed at least 300 trials in an hour.

### Electrophysiological Recordings and Spike Sorting

To record neural activity with a high degree of temporal resolution, we implanted 16-channel microdrive-driven array recording optrodes (2×8 electrode grid, 35µm diameter/electrode, Innovative Neurophysiology) into either the anterior dorsomedial striatum (A-DMS; AP: +1.15 mm, ML: +1.35 mm, DV: −2.8 mm), the posterior dorsomedial striatum (P-DMS; AP: −0.25 mm, ML: +1.9 mm, DV: −2.7 mm) or the Substantia Nigra *pars reticulata* (SNr; AP: -3.5mm, ML:+1.5mm, DV: -4.0mm). Twenty-four hours prior to recording, electrodes were advanced in 100 µm increments to fully sample the dorsoventral (D-V) extent of the DMS (five positions per mouse). We used semi-automated spike sorting software (Kilosort v2.5) to isolate single-unit activity and generate putative spike clusters. Each cluster was then manually validated for single-unit isolation using Phy2.0.

### Analysis of *In-vivo* Electrophysiological Recordings

To account for intrasession and intra-animal variability in trial execution, we defined five epochs in the task based on the median length of time it took all animals to perform each part of the trial: 1.5 seconds from start of the ITI until initiation,1 second from initiation until the choice, 1 second from the choice until returning to the reward port, 0.6 seconds from reward port entry until reward delivery, and 0.9 seconds for reward consumption (Fig.2B). Those epochs were aligned across trial and animal by implementing the time-warping algorithm, which integrates the variability of the epochs across trials to generate a fixed time for trials of varying length (‘warped’).

We used custom MATLAB scripts to calculate the firing rate of isolated units in windows of 10ms for each trial in warped time. To determine the percentage of modulated units in each epoch, we performed a receiver operating characteristic (ROC) curve analysis to determine whether the spike frequency in each time bin differed from that of the baseline time taken at the beginning of the ITI. To calculate spike frequency, we used a 500ms window at the baseline and a sliding window of 50ms to compare the spike frequency of each unit at each time point compared to the baseline. To calculate the significance of the area under the curve (auROC) value, we compared this value against a null distribution generated via 10,000 permutations of the baseline and time window distributions. Only significant auROC values (p < 0.01) were used to compare across different behavioral parameters. To examine population recruitment, we used the above sliding window auROC approach either on unsorted data (Fig.S2A,B; Fig.S13A), ‘extinction’ trials (Fig.S2A,B; Fig.S13A), current trial choice (Fig.2J-M; Fig.6J), Q_contra_ (Fig.3A-D; Fig6K), Q* (Fig.3H,I; Fig.6L), or Q_ipsi_ (Fig.S3H,I; Fig.S13E). To compare recruitment according to Q-values, we fit a linear model with the number of units recruited within each trial epoch for different Q-values using Prism’s simple linear regression function.

#### Principal Component Analysis (PCA)

The PCA was implemented to understand the neural dynamics of the A-DMS or P-DMS population responses. We used the mean firing rate of the time-warped traces, then applied the *pca* function from MATLAB to calculate the first three principal components. The population activity was projected onto the first three principal component axes to describe the population’s activity trajectories across the task’s trial epochs.

#### Linear regression

To address whether an individual neurons’ firing rate had a relationship with Q-values, we performed a linear regression model on the trial-by-trial firing rates (Figure S3D,E) with Q_contra_ and Q_ipsi_ as predictors. We z-scored the firing rates and then used the *regress* function from MATLAB to determine whether there was a relationship between firing rates and Q values.

### Optogenetics

To evaluate the functional role of the A-DMS and the P-DMS, we used a halorhodopsin virus to locally inhibit neurons in these areas as animals performed the task. Mice were bilaterally injected in the A-DMS or P-DMS with 150 nl of either AAV5 hSyn-eNpHR3.0-eYFP-WPRE-hGH (n=12 NpHR mice per brain area) or rAAV5 hSyn-EYFP (n=8 control mice per brain area). Custom-made optic cannulas (Thorlabs, FT200EMP, SFLC230) were bilaterally implanted dorsal to the injection sites so that light the opsin-activating light (532nm, 4-5mW measured at the fiber tip) could be delivered into the striatum. Animals were given 14 days of recovery before starting food deprivation and behavioral training. After animals were trained and reached the criteria for task performance, they performed a series of behavioral sessions with pre- or post-choice inhibition performed unilaterally or bilaterally. Within each session, opsin-activating light was active on every trial in half of the blocks. We set 400 trials, 65% accuracy, and unbiased choice (no more than 2:1 between sides) as criteria for inclusion of any behavioral session into the final dataset. We performed histology on animals to confirm virus targeting and fiber placement. There were two A-DMS implanted animals where viral expression was confirmed, but precise fiber placement could not be confirmed due to tissue damage.

### Modeling of animal choice

We used the following logistic regression model to analyze the extent to which reward and choice history contribute to animals’ choice on a trial-by trial basis:

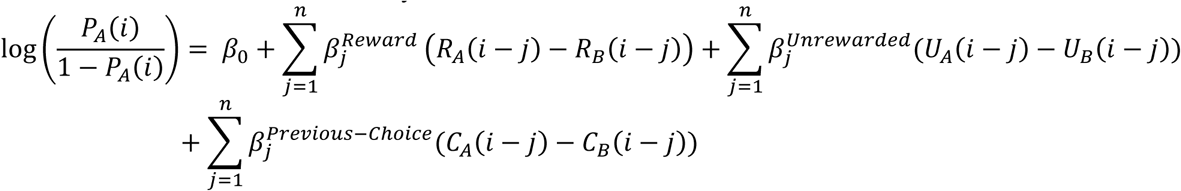

where *P*_*A*_(*i*) is the probability of selecting choice *A* on trial *i*. *R*_*A*_ and *R*_*B*_ are variables that represent whether a reward was given (*R*=1 for 12µL reward; *R*=0.333 for 4 µL reward) or not given (*R*=0) in port *A* or port *B* respectively. *U*_*A*_ and *U*_*B*_ are variables that represent whether a choice was unrewarded (*U*=1) or not unrewarded (*U*=0) in port *A* or port *B* respectively. *C*_*A*_ and *C*_*B*_ are variables that represent whether a choice was repeated (*C*=1) or not repeated (*C*=0) in port *A* or port *B* respectively. The variable *n* represents the number of trials that were included in the model fits (*n*=3 here). The regression coefficient *β*_0_ represents the bias of the animal. The regression coefficients, 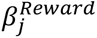, 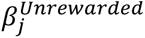, and 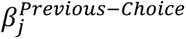, represent the extent to which prior rewarded trials, unrewarded trials, and choices impact the current choice. We used MATLAB’s *glmfit* function to estimate the corresponding coefficients.

We adapted a Q-learning Reinforcement Learning Model with three parameters to fit the behavioral data produced by the volume comparison task. Mouse choice and outcome history were the primary inputs of the model. The values of the choice alternatives were initiated at 0 and updated as follows. For the chosen port,

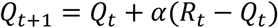

where *Q*_*t*_ is the value of the action taken on trial t of each choice and *R*_*t*_ is the actual reward received in trial t. The parameter α is the learning rate, which controls the degree to which the outcome of a certain trial is used to estimate value of choices on future trials. The decision process mapping the Q-values to the probability of choosing one port over the other was modeled with a softmax rule:

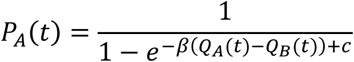

where *P*_*A*_(*t*) is the probability of choosing port *A* on trial t. Here, *β* is the inverse temperature parameter which controls the degree to which choices are biased by differences in estimated value. High values for *β* indicate that mice more readily exploit differences in action values between the alternatives, while lower values suggest that mice exhibit more exploratory behavior. Here, *c* is the bias term, which captures the degree to which value-based choice is affected by bias. A positive bias term would bias the animal toward option *A*, whereas a negative bias term would bias the animal toward option *B*. To fit this model to our choice data, we used the *fmincon* function in MATLAB to minimize the negative log likelihood of models using our parameters (α, β, *c*). For optogenetic experiments, our unique task and manipulation protocol allowed us to fit separate parameters for light-off trials (α, β, *c*) and light-on trials (α’, β’, *c*’). We used a *k-folds* strategy to alternate training data with different subsets of testing data. By taking the average of the various fits, we were able to reduce biases from training data and improve model likelihood.

### Open Field Video Tracking

To test for a relationship between optogenetically induced choice bias and movement, we performed inhibition as mice explored an open field. Animals were placed in an open field chamber (12”L by 15”W by 12”H, custom made from clear plastic) housed inside of a sound-attenuating cabinet and given 1 minute to habituate. Afterwards, we delivered intermittent opsin-activating light (532nm, 4-5mW) for 1.5 seconds followed by 13.5 seconds without light. A total of 80 intervals of light activation were recorded for each mouse. We recorded open field sessions using a webcam (Brio, Logitech) placed underneath the open field chamber. Video recordings were taken at 10fps and TTLs for light activation were recorded using an RZ5 processor (Tucker Davis Technologies, Synapse Software). We used *Deeplabcut*^53^ (Version 2.2.0.6) to perform tracking of the nose, body, and tail base coordinates as mice explored the open field. We used the distance traveled of the tail base over 5 frames to calculate the animal’s velocity. We used the cosine rule to calculate body angle using the distances between our tracking points. We took a one second window around the light turning on or off and measured changes in velocity and body angle to infer whether the optogenetic inhibition was having any purely motor effect (Fig.S6).

### Anatomical Tracing

To address whether the A-DMS and P-DMS have a common output, we used a transynaptic strategies injecting AAV1-hSyn-cre and AAV1-hSyn-Flp (470nL) into the A-DMS (AP:+1.2mm, ML:+1.4mm, DV:-2.7mm ) or P-DMS (AP:+0.1mm, ML:+1.9mm, DV:-2.7mm ), respectively. These injections were counterbalanced across animals such that half received AAV1-cre into the A-DMS, and half received AAV1-FLP into the A-DMS; the other virus was injected into the P-DMS. Striatal injections were combined with 30 nL of CTB647 to confirm the site of the injection. We then injected a combination of viruses (AAV5-flex-mCherry & AAV5-fDIO-EYFP) into the SNr (AP:-3.5mm, ML:+1.5mm, DV:-4.2mm). After one month of expression, mice were anesthetized with pentobarbital then perfused with PBS followed by 4% paraformaldehyde (PFA) in PBS. Brains were post-fixed overnight at 4°C in 4% PFA, after which PFA was removed and replaced with PBS. A vibratome was used to cut 50-μm sections and sections were mounted on slides and coverslipped with VectaMount mounting media (Vector Laboratories).

Images were captured at 10X magnification using an epifluorescent microscope (Leica DM6) and stitched into compilations spanning the entire slide. We used separate channels to capture the mCherry, eYFP, and CTB647 expression. We used a custom script in ImageJ to isolate the numbers and locations of neurons labeled by mCherry and eYFP across a range of SNr coordinates (AP: -2.755:-3.855), then used a custom MATLAB script to overlay these coordinates and calculate local density across the SNr (Fig.6C,D).

### Quantification and Statistical Analysis

All data were analyzed with Prism 10.4 and custom MATLAB or Python code, available upon request. Repeated Measures ANOVAs, One-way ANOVAs, and non-parametric Wilcoxon matched pairs rank sum test were performed using Prism 10.4’s built-in functions to compare grouped data. We used Prism 10.4’s function to perform Fisher’s exact test and the McNemar test to measure significance of 2×2 contingency tables where appropriate. We used Prism 10.4’s regression function to perform linear regressions of neuronal recruitment across Q- and Q*-values. We used MATLAB’s *glmfit* function to perform regression of current trial choice according to prior choices and reward outcomes. We also used Prism 10.4’s function to perform Kruskal-Wallis tests for non-parametric comparisons of different groups where appropriate.

## Acknowledgements

This research was supported by grants NIMH R01 MH118369 (M.V.F), NIMH R01 MH136354 (M.V.F), NINDS R01 NS117061 (M.V.F) and NIMH F31 MH125542 (L.V.C). We thank Alessandro Jean-Louis and Marcella Soewignjo for excellent technical assistance. We also thank Sarah Ferrigno and Evan Illiakis for careful reading of the manuscript.

## Author Contributions

L.V.C., E.D.H., and M.V.F. conceptualized the project. L.V.C. performed and analyzed all the optogenetics experiments and the behavior modeling. E.D.H. performed and analyzed all neural recordings. M.G. and W.T. performed all anatomical tracing experiments. L.V.C., E.D.H., and M.V.F. wrote the manuscript.

## Notes

### Competing Interest Statement

The authors have declared no competing interest.

