## Supplemental Figures for "Temporally and Functionally Distinct Contributions to Value Based Choice Along the Anterior-Posterior Dorsomedial Striatal Axis"

SUPPLEMENTARY FIGURES

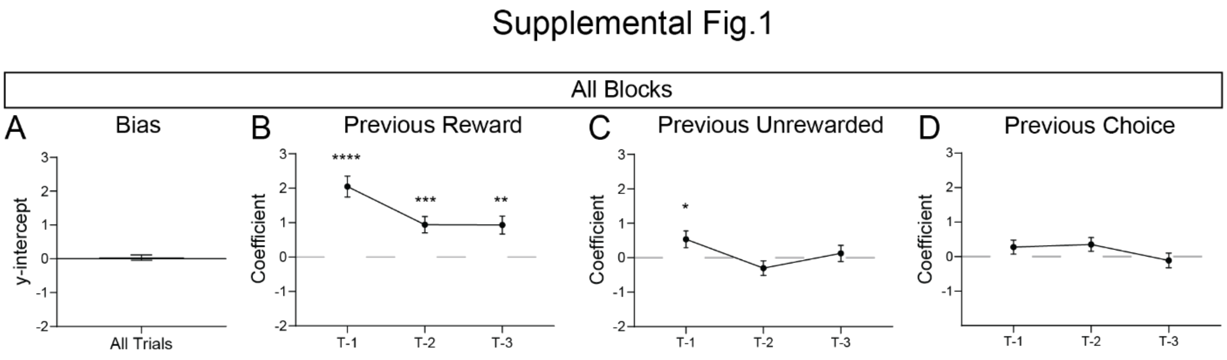


**Supplementary Figure 1. Logistic regression on volume contrast task performance indicates multi-trial integration of value**

(A) Mean logistic regression y-intercept value, which captures bias for making a choice to the right

(B) Coefficients for rewarded trials for previous three trials show integration of reward for previous three trials to current choice (*n*=27; one sample t-test, T-1: t=6.748, ****P < 0.0001, T-2: t=3.955, ***P = 0.0005, T-3: t=3.574, **P = 0.0014).

(C) Coefficient for unrewarded trials on previous three trials show a modest contribution for the previous trial to current choice (*n*=27; one sample t-test, T-1: t=2.193, *P = 0.0371, T-2: t=1.434, P = 0.1632, T-3: t=0.5320, P = 0.5992).

(D) Coefficient for previous choice shows that there is no contribution of prior choice to current choice (*n*=27; one sample t-test, T-1: t=1.360, P = 0.1850, T-2: t=1.787, P = 0.0853, T-3: t=0.5249, P = 0.6041).

Supplemental Fig.2

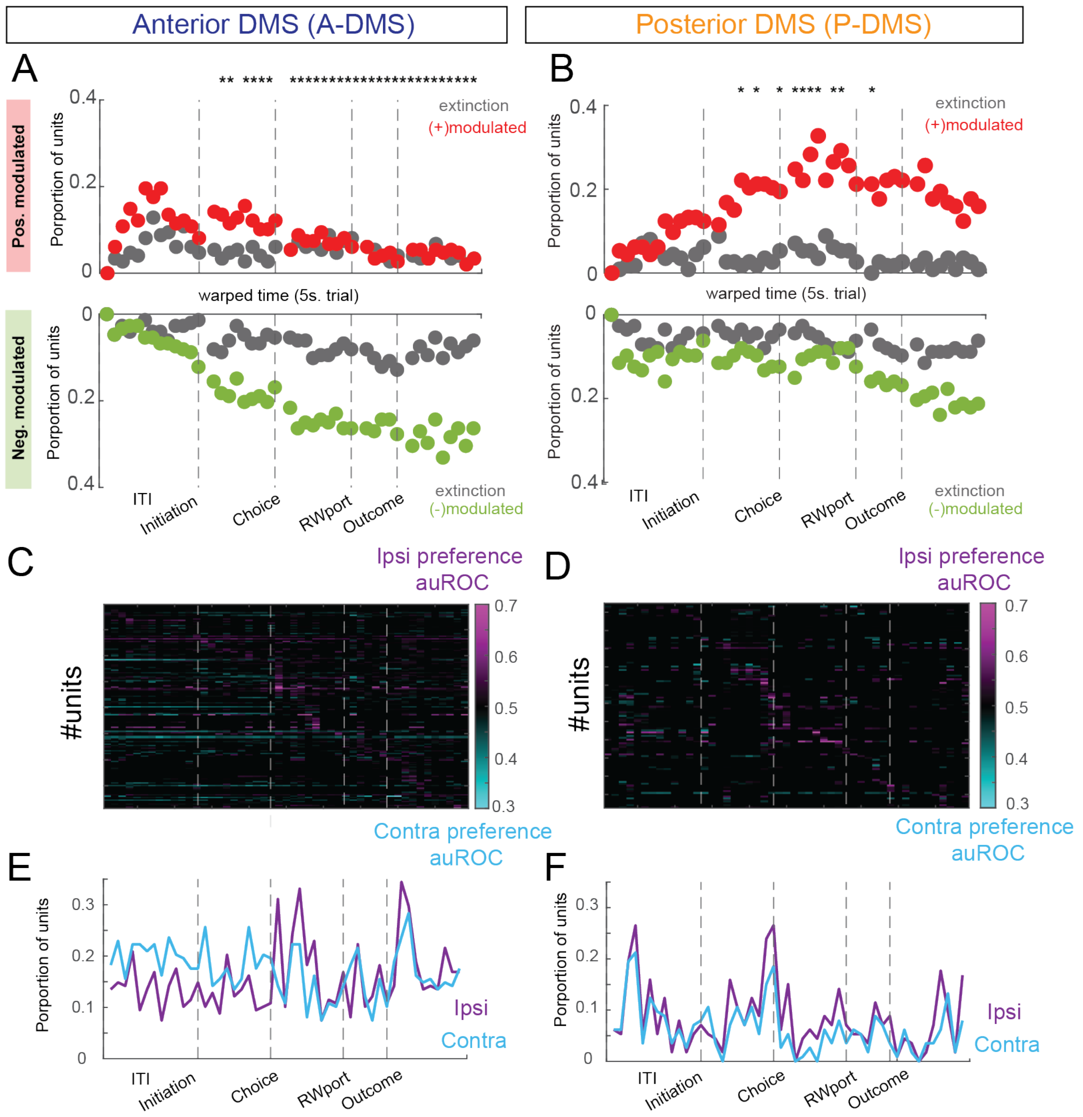


**Supplementary Figure 2. Modulation of activity in the A- and P-DMS for extinction trials and contralateral vs ipsilateral choice**

(A) Positive (top) and negative (bottom) modulation of A-DMS units for trials 1-25 (red) and extinction trials (26-30, gray) show a reduction in modulation for extinction trials in the A-DMS. * $p<0.001$ McNemar's test

(B) Positive (top) and negative (bottom) modulation of P-DMS units for trials 1-25 (green) and extinction trials (gray) shows there is a reduction in modulation for extinction trials in the P-DMS. * $p<0.001$ McNemar's test

(C) Heat map showing individual unit choice tuning for A-DMS neurons as measured by auROC for ipsilateral vs contralateral choice firing rates. Purple and blue represent ipsi and contra preference, respectively.

(D) Same as C, but for P-DMS neurons.

(E) Proportion of A-DMS neurons with ipsilateral or contralateral choice tuning (auROC ≠ 0.5) throughout the trial.

(F) Same as E, but for P-DMS.

### Supplemental Fig.3

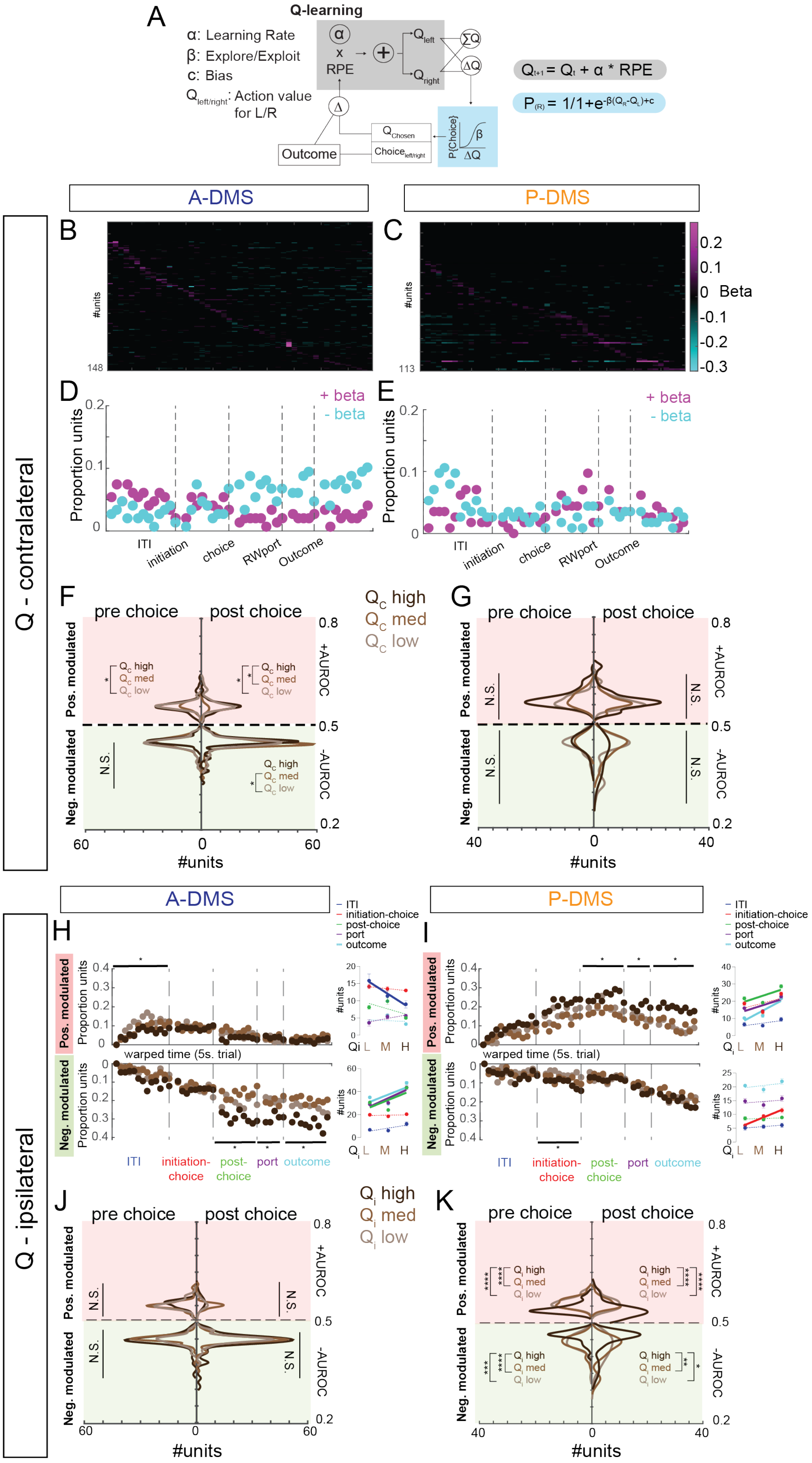

**Supplementary Figure 3. Modulation of activity in the A- and P-DMS according to values of contralateral and ipsilateral choices**

(A) Q-Learning model with parameters for learning rate, explore/exploit tendency, and choice bias. Choice modeled using softmax function (blue) and formula for Q value updating (grey).

(B) Heat map of the beta regression parameters for firing rates of neurons in the A-DMS against the $Q_{contra}$ (ranked by temporal location of peak beta).

(C) Same as B, but for the P-DMS.

(D and E) Fewer than 10% of neurons showed firing rate changes as a function of value in either the A-DMS (D) or the P-DMS (E).

(F) Plot of the number of modulated units (x-axis) and their respective modulation indices (auROC, y-axis) for three $Q_{contra}$ values, separated by positive (pink) and negative (green) modulation as well as by pre- or post-choice (left versus right half). auROC values for individual units did not demonstrate positive modulation of A-DMS units for higher $Q_{contra}$ values in the pre-choice period (Top Left: Kruskal-Wallis test with Dunn's test for multiple comparisons, $H_2 = 1.267$, P=0.5306). However, we did observe positive modulation of units in the post-choice period (Top Right: Kruskal-Wallis test with Dunn's test for multiple comparisons, $H_2 = 7.264$, *P=0.0265; *Post-hoc* comparisons: **p<0.05*). Individual units also scaled their activity towards lower auROC values as a function of value in the A-DMS for the pre-choice period (Bottom Left: Kruskal-Wallis test with Dunn's test for multiple comparisons, $H_2 = 6.528$, *P=0.0382; *Post-hoc* comparisons: *p<0.05) and the post-choice period (Bottom Right: Kruskal-Wallis test, $H_2 = 8.118$, *P=0.0173; *Post-hoc* comparisons: *p<0.05).

(G) Same as F, but for P-DMS neurons. Individual units did not scale activity towards higher auROC values as a function of value for positive modulation in the pre-choice period (Top Left: Kruskal-Wallis test with Dunn's test for multiple comparisons, $H_2 = 4.876$, P=0.0873) or the post-choice period (Top Right: Kruskal-Wallis test with Dunn's test, $H_2 = 0.6732$, P=0.7142). Similarly, units did not scale their activity towards lower auROC values as a function of value for negative modulation in the pre-choice period (Bottom Left: Kruskal-Wallis test, $H_2 = 0.2102$, P=0.9002) or the post-choice period (Bottom Right: Kruskal-Wallis test, $H_2 = 3.139$, P=0.2081).

(H) Top: changes in the proportion of positively modulated units at different values for ipsilateral choice in the A-DMS. Positive modulation inversely correlates with value in the ITI window for A-DMS. Bottom: Negative modulation robustly scales with value from the post-choice window through the outcome window in the A-DMS. Insets show linear regression for number of units modulated within each window according to value.

(I) Same as H, but for P-DMS. Top: positive modulation in the P-DMS scales with $Q_{ipsi}$ from the post-choice window through the outcome window in the P-DMS. Bottom: Negative modulation in the P-DMS weakly scales with value in the initiation-choice period only.

(J) Same as F, but for $Q_{ipsi}$. Individual units for not scale activity towards higher auROC values as a function of value for positive modulation in the pre-choice period (Top Left: Kruskal-Wallis test, $H_2 = 1.798$, P=0.4069) or the post-choice period (Top Right: Kruskal-Wallis test, $H_2 = 5.742$, P=0.0567). Similarly, units did not scale their activity towards lower auROC values as a function of value for negative modulation in the pre-choice period (Bottom Left: Kruskal-Wallis test, $H_2 = 5.366$, P=0.0684) or the post-choice period (Bottom Right: Kruskal-Wallis test, $H_2 = 2.464$, P=0.2917).

(K) Same as J, but for P-DMS neurons. auROC values for individual units reveal that there is increased positive modulation of P-DMS units for lower $Q_{ipsi}$ values in both the pre-choice period (Top Left: Kruskal-Wallis test with Dunn's test for multiple comparisons, $H_2 = 54.75$, ****P<0.0001; *Post-hoc* comparisons: *****p<0.0001*) and the post-choice period (Top Right: Kruskal-Wallis test with Dunn's test for multiple comparisons, $H_2 = 49.56$, ****P<0.0001; *Post-hoc* comparisons: *****p<0.0001*). Individual units also scaled their activity towards increased negative modulation as a function of value in the P-DMS for the pre-choice period (Bottom Left: Kruskal-Wallis test with Dunn's test for multiple comparisons, $H_2 = 27.25$, ****P<0.0001; *Post-hoc* comparisons: ***p<0.001, ****p<0.0001) and the post-choice period (Bottom Right: Kruskal-Wallis test with Dunn's test for multiple comparisons, $H_2 = 12.11$, **P=0.0024; *Post-hoc* comparisons: *p<0.05, **p<0.01).

Supplemental Fig.4

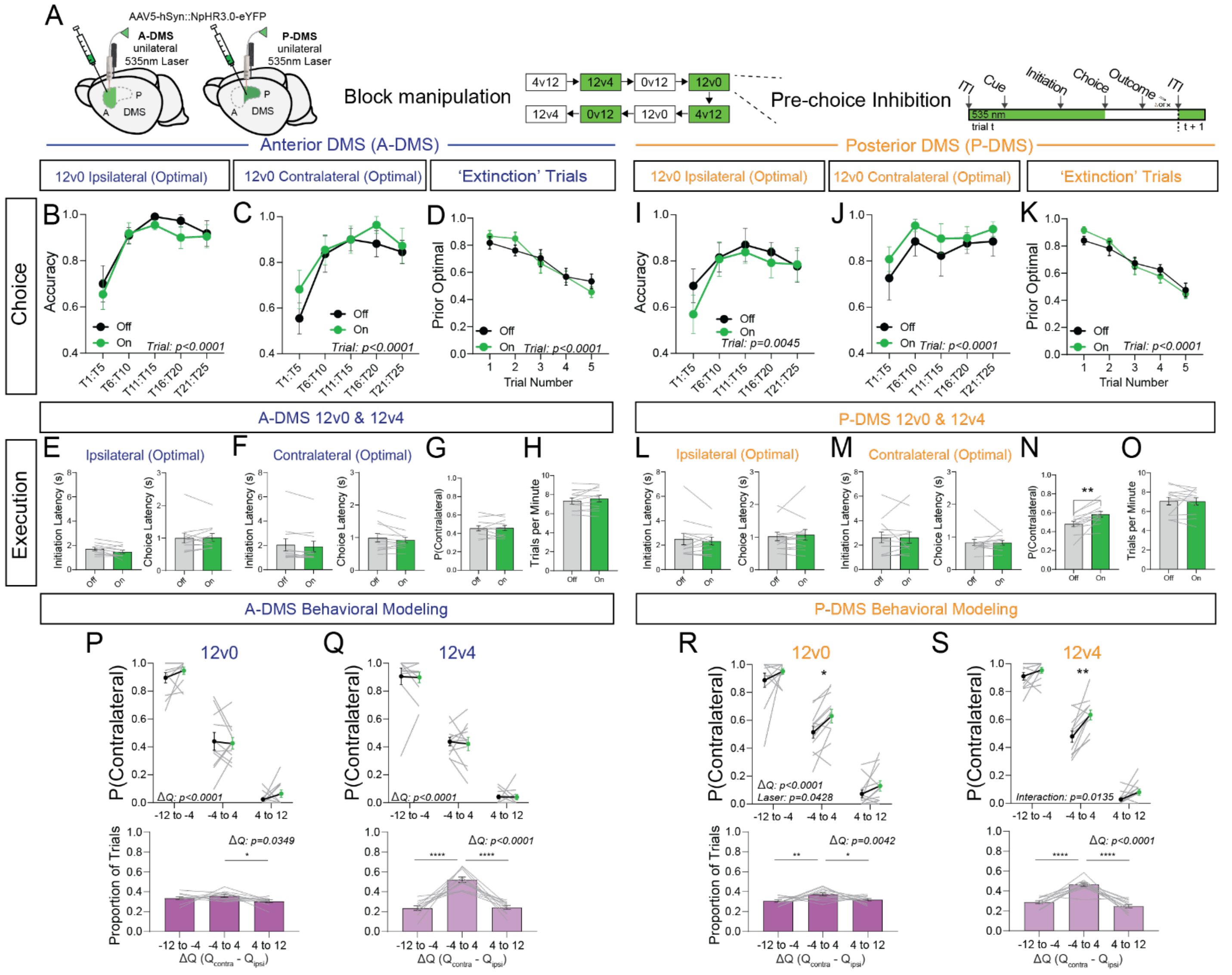


**Supplementary Figure 4. Pre-choice manipulation in P-DMS, but not A-DMS, impacts choice**

(A) Experiment schematic. Unilateral activation of NpHR3.0 in the A- or P-DMS in 50% of blocks. Every trial in a manipulation block gets 532nm light delivery during the pre-choice period.

(B) Proportion of optimal choices made by A-DMS:NpHR animals in 12v0 blocks when the laser is ipsilateral or contralateral to the optimal choice and laser is off (black) or the laser is on (green) (*n*=11; RM-two-way ANOVA, trial: $F_{2,23}$ = 14.57, ****P < 0.0001, Laser: $F_{1,10}$ = 0.6391, P = 0.4426, interaction: $F_{2,20}$ = 0.2711, P = 0.7639).

(C) Same as B, but for 12v0 blocks when the laser is contralateral to the optimal choice (*n*=11; RM-two-way ANOVA, trial: $F_{3,30}$ = 10.60, ****P < 0.0001, Laser: $F_{1,10}$ = 1.441, P = 0.2577, interaction: $F_{3,26}$ = 0.4490, P = 0.6594).

(D) Proportion of choices made by A-DMS:NpHR to the prior optimal side in 'extinction' trials when the laser is on (green) or off (black) (*n*=11; RM-two-way ANOVA, trial: $F_{3,28}$ = 17.27, ****P < 0.0001, Laser: $F_{1,10}$ = 0.01975, P = 0.8910, interaction: $F_{2,24}$ = 1.511, P = 0.2399).

(E) Initiation and choice latencies (left and right, respectively) for A-DMS:NpHR animals making optimal choices ipsilateral to the laser manipulation side when the laser was off (gray) or on (green) (*n*=11; Wilcoxon matched pairs rank sum test, Initiation Latency: P = 0.0830, Choice Latency: P = 0.8311).

(F) Same as in E, but for animals making optimal choices contralateral to the laser manipulation side (*n*=11; Wilcoxon matched pairs rank sum test, Initiation Latency: P = 0.1230, Choice Latency: P = 0.5771).

(G) Probability that A-DMS:NpHR animals made choices to the contralateral side when the laser was off (gray) or on (green) (*n*=11; Wilcoxon matched pairs rank sum test, P(Contralateral): P = 0.8457).

(H) Average number of trials completed per minute for A-DMS:NpHR animals when the laser was off (gray) or on (green) (*n*=11; Wilcoxon matched pairs rank sum test, Trial Rate: P = 0.2783).

(I) Proportion of optimal choices made by P-DMS:NpHR animals in 12v0 blocks when the laser is ipsilateral to the optimal choice and laser is off (black) or the laser is on (green) (*n*=12; RM-two-way ANOVA, trial: $F_{2,26}$ = 6.238, **P = 0.0045, Laser: $F_{1,11}$ = 1.303, P = 0.2780, interaction: $F_{2,21}$ = 0.3272, P = 0.7168).

(J) Same as I, but for 12v0 blocks when the laser is contralateral to the optimal choice (*n*=12; RM-two-way ANOVA, trial: $F_{2,25}$ = 3.031, P = 0.0604, Laser: $F_{1,11}$ = 1.179, P = 0.3007, interaction: $F_{2,24}$ = 0.1190, P = 0.9045).

(K) Proportion of choices made by P-DMS:NpHR animals to the prior optimal side in 'extinction' trials when the laser is on (green) or off (black) (*n*=12; RM-two-way ANOVA, trial: $F_{3,26}$ = 34.38, ****P < 0.0001, Laser: $F_{1,11}$ = 0.02466, P = 0.8781, interaction: $F_{3,36}$ = 1.013, P = 0.4028).

(L) Initiation and choice latencies (left and right, respectively) for P-DMS:NpHR animals making optimal choices ipsilateral to the laser manipulation side when the laser was off (gray) or on (green) (*n*=12; Wilcoxon matched pairs rank sum test, Initiation Latency: P = 0.5693, Choice Latency: P = 0.9097).

(M) Same as in L, but for animals making optimal choices contralateral to the laser manipulation side (*n*=12; Wilcoxon matched pairs rank sum test, Initiation Latency: P > 0.9999, Choice Latency: P = 0.9697).

(N) Probability that P-DMS:NpHR animals made choices to the contralateral side when the laser was off (gray) or on (green) (*n*=12; Wilcoxon matched pairs rank sum test, P(Contralateral): ***P = 0.0010).

(O) Average number of trials completed per minute for P-DMS:NpHR animals when the laser was off (gray) or on (green) (*n*=12; Wilcoxon matched pairs rank sum test, Trial Rate: P = 0.4238).

(P) Top: No change in probability of A-DMS:NpHR animals to make a contralateral choice at various ΔQ ($Q_{ipsi} - Q_{Contra}$) values in 12v0 blocks when the laser was off (black) or on (green). Bottom: Proportion of trials within each ΔQ bin are shown (bottom) (*n*=11; RM-two-way ANOVA (top), Delta Q: $F_{1,13}$ = 338.9, ****P < 0.0001, Laser: $F_{1,10}$ = 1.264, P = 0.2872, Interaction: $F_{2,20}$ = 0.6723, P = 0.5163. One-way ANOVA with Tukey's correction for multiple comparisons, Bin Proportions: F = 3.760, *P = 0.0349; *Post-hoc* comparisons: **p* < 0.05).

(Q) Same as in P, but for trials in 12v4 blocks (*n*=11; RM-two-way ANOVA (top), Delta Q: $F_{2,17}$ = 213.6, ****P < 0.0001, Laser: $F_{1,10}$ = 0.1240, P = 0.7320, Interaction: $F_{2,16}$ = 0.05034, P = 0.9239. One-way ANOVA with Tukey's correction for multiple comparisons (bottom), Bin Proportions: F = 47.77, *P < 0.0001; *Post-hoc* comparisons: ****p < 0.0001).

(R) Increased probability of P-DMS:NpHR animals to make a contralateral choice at various ΔQ ($Q_{ipsi} - Q_{Contra}$) values in 12v0 blocks when the laser was off (black) or on (green) (top). Proportion of trials within each ΔQ bin are shown (bottom) (*n*=12; RM-two-way ANOVA (top), Delta Q: $F_{2,20}$ = 301.3, ****P < 0.0001, Laser: $F_{1,11}$ = 5.241, *P = 0.0428, Interaction: $F_{2,19}$ = 0.9256, P = 0.4000. One-way ANOVA with Tukey's correction for multiple comparisons (bottom), Bin Proportions: F = 6.497, **P = 0.0042; *Post-hoc* comparisons: **p* < 0.05, ***p* < 0.01).

(S) Same as in P, but for trials in 12v4 blocks (*n*=12; RM-two-way ANOVA (top), Delta Q: $F_{1,14}$ = 411.0, ****P < 0.0001, Laser: $F_{1,11}$ = 16.92, **P = 0.0017, Interaction: $F_{2,21}$ = 5.367, *P = 0.0137. One-way ANOVA with Tukey's correction for multiple comparisons (bottom), Bin Proportions: F = 47.77, *P < 0.0001; *Post-hoc* comparisons: ***p* < 0.01, ****p < 0.0001).

### Supplemental Fig.5

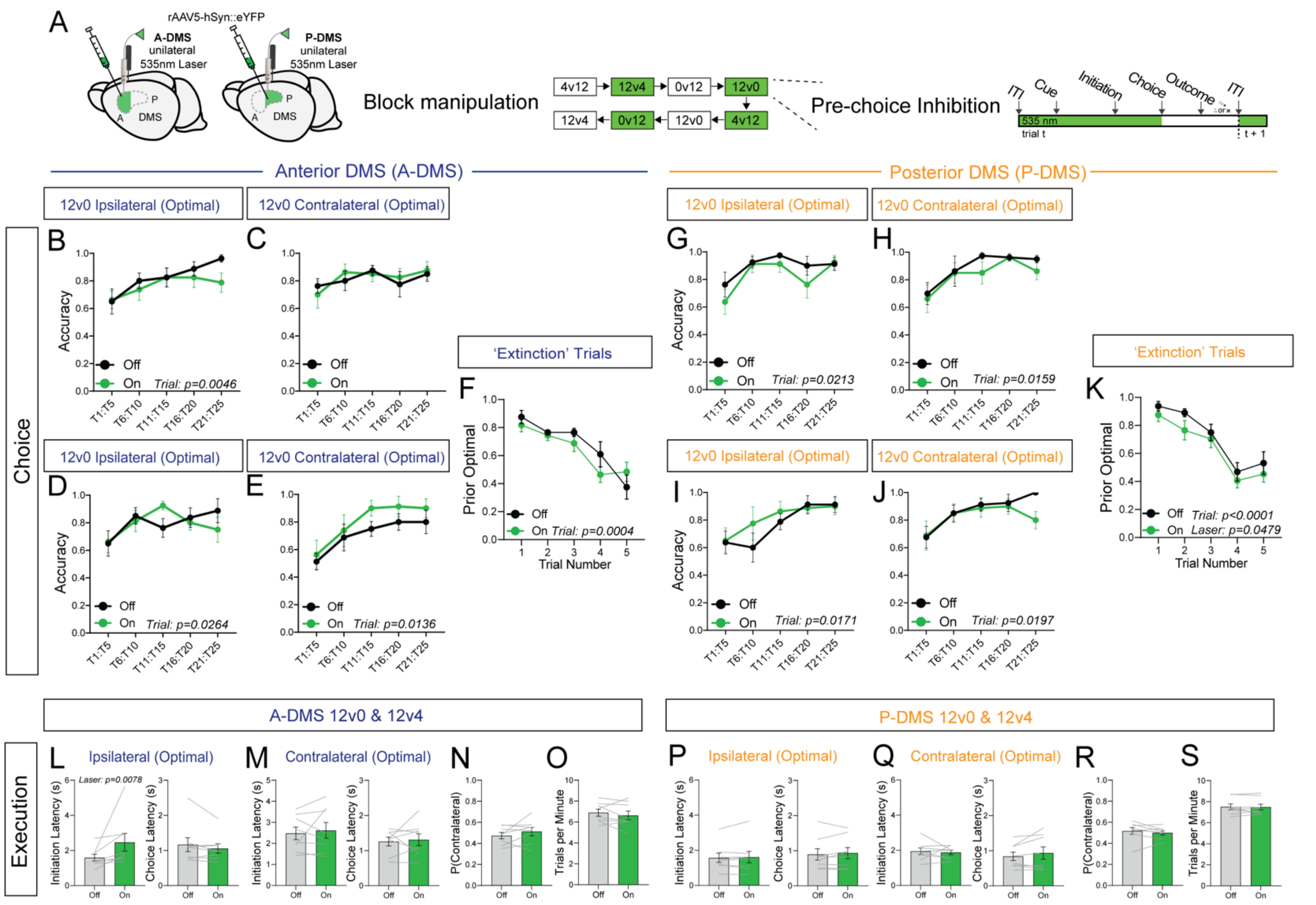


**Supplementary Figure 5. GFP controls for unilateral pre-choice inhibition**

(A) Experiment schematic. Unilateral activation of GFP in the A- or P-DMS in 50% of blocks. Every trial in a manipulation block gets 532nm light delivery during the pre-choice period.

(B) Proportion of optimal choices made by A-DMS:GFP animals in 12v0 blocks when the laser is ipsilateral to the optimal choice and the laser is off (black) or the laser is on (green) (*n*=8; RM-two-way ANOVA, trial: $F_{2,15}$ = 7.475, **P = 0.0046, laser: $F_{1,7}$ = 0.4777, P = 0.5177, interaction: $F_{2,25}$ = 1.162, P = 0.3465).

(C) Same as B, but for 12v0 blocks when the laser is contralateral to the optimal choice (n=8; RM-two-way ANOVA, trial: $F_{3,18}$ = 1.665, P = 0.2133, laser: $F_{1,7}$ = 0.1045, P = 0.7560, interaction: $F_{2,23}$ = 0.4024, P = 0.6980).

(D) Proportion of optimal choices made by A-DMS:GFP animals in 12v4 blocks when the laser is ipsilateral to the optimal choice and the laser is off (black) or the laser is on (green) (*n*=8; RM-two-way ANOVA, trial: $F_{2,17}$ = 4.244, *P = 0.0264, laser: $F_{1,7}$ = 0.01283, P = 0.9130, interaction: $F_{3,20}$ = 2.074, P = 0.1374)

(E) Same as D, but for 12v4 blocks when the laser is contralateral to the optimal choice (n=8; RM-two-way ANOVA, trial: $F_{2,15}$ = 5.718, *P = 0.0136, laser: $F_{1,7}$ = 4.718, P = 0.0664, interaction: $F_{3,20}$ = 0.1966, P = 0.7672)

(F) Proportion of choices made by A-DMS:GFP to the prior optimal side in 'extinction' trials when the laser is on (green) or off (black) (*n*=8; RM-two-way ANOVA, trial: $F_{2,14}$ = 14.15, ***P = 0.0004, laser: $F_{1,7}$ = 0.7902, P = 0.4035, interaction: $F_{2,13}$ = 2.018, P = 0.1736)

(G) Proportion of optimal choices made by P-DMS:GFP animals in 12v0 blocks when the laser is ipsilateral to the optimal choice and the laser is off (black) or the laser is on (green) (*n*=8; RM-two-way ANOVA, trial: $F_{2,14}$ = 4.850, *P = 0.0213, laser: $F_{1,7}$ = 2.015, P = 0.1987, interaction: $F_{2,14}$ = 1.294, P = 0.3051)

(H) Same as G, but for 12v0 blocks when the laser is contralateral to the optimal choice (*n*=8; RM-two-way ANOVA, trial: $F_{2,11}$ = 6.815, *P = 0.0159, laser: $F_{1,7}$ = 1.184, P = 0.3126, interaction: $F_{3,19}$ = 0.4364, P = 0.7107)

(I) Proportion of optimal choices made by P-DMS:GFP animals in 12v4 blocks when the laser is ipsilateral to the optimal choice and the laser is off (black) or the laser is on (green) (*n*=8; RM-two-way ANOVA, trial: $F_{2,11}$ = 6.745, *P = 0.0171, laser: $F_{1,7}$ = 0.9861, P = 0.3538, interaction: $F_{2,11}$ = 1.535, P = 0.2547)

(J) Same as H, but for 12v4 blocks when the laser is contralateral to the optimal choice (n=8; RM-two-way ANOVA, trial: $F_{2,12}$ = 5.830, *P = 0.0197, laser: $F_{1,7}$ = 1.629, P = 0.2425, interaction: $F_{3,18}$ = 1.702, P = 0.2048)

(K) Proportion of choices made by P-DMS:GFP animals to the prior optimal side in 'extinction' trials when the laser is on (green) or off (black) (*n*=8; RM-two-way ANOVA, trial: $F_{3,18}$ = 31.97, ****P < 0.0001, laser: $F_{1,7}$ = 5.727, *P = 0.0479, interaction: $F_{3,18}$ = 0.1795, P = 0.8816)

(L) Initiation and choice latencies (left and right, respectively) for A-DMS:GFP animals making choices ipsilateral to the laser manipulation side when the laser was off (gray) or on (green) (*n*=8; Wilcoxon matched pairs rank sum test, Initiation Latency: **P = 0.0078, Choice Latency: P = 0.1953).

(M) Same as in L, but for choices made when the optimal side was contralateral to the laser manipulation side (n=8; Wilcoxon matched pairs rank sum test, Initiation Latency: P = 0.9453, Choice Latency: P = 0.8438).

(N) Probability that A-DMS:GFP animals made choices to the contralateral side when the laser was off (gray) or on (green) (*n*=8; Wilcoxon matched pairs rank sum test, P(Contralateral): P = 0.3828).

(O) Average number of trials completed per minute for A-DMS:GFP animals when the laser was off (gray) or on (green) (*n*=8; Wilcoxon matched pairs rank sum test, Trial Rate: P = 0.3828).

(P) Initiation and choice latencies (left and right, respectively) for P-DMS:GFP animals making choices ipsilateral to the laser manipulation side when the laser was off (gray) or on (green) (*n*=8; Wilcoxon matched pairs rank sum test, Initiation Latency: P = 0.9453, Choice Latency: P = 0.8438).

(Q) Same as P, but for choices made when the optimal side was contralateral to the laser manipulation side (n=8; Wilcoxon matched pairs rank sum test, Initiation Latency: P = 0.6406, Choice Latency: P = 0.3828).

(N) Probability that P-DMS:GFP animals made choices to the contralateral side when the laser was off (gray) or on (green) (n=8; Wilcoxon matched pairs rank sum test, P(Contralateral): P = 0.3828).

(O) Average number of trials completed per minute for P-DMS:GFP animals when the laser was off (gray) or on (green) (n=8; Wilcoxon matched pairs rank sum test, Trial Rate: P > 0.9999).

Supplemental Fig.6

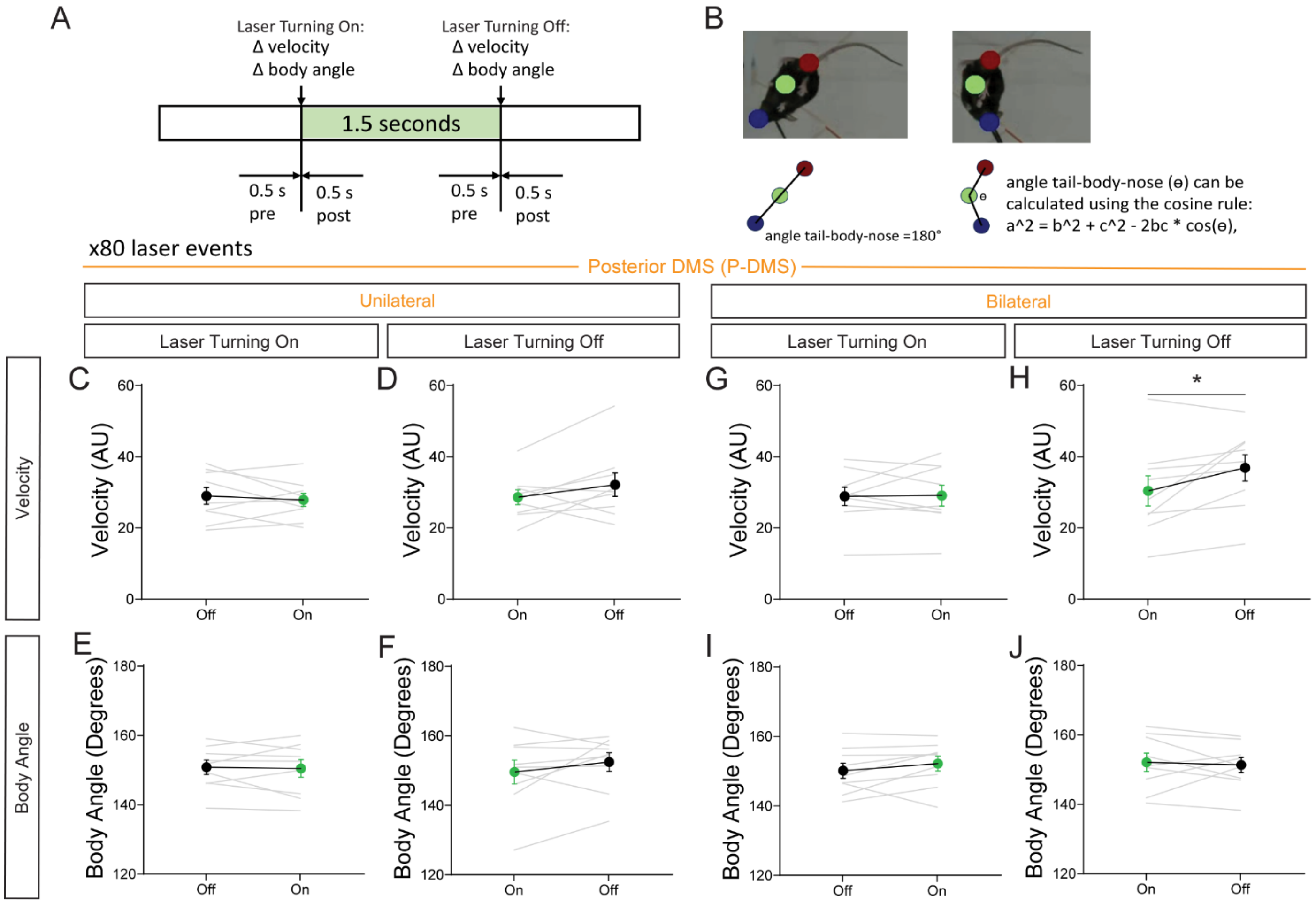


**Supplementary Figure 6. Unilateral P-DMS inhibition does not induce motor effects in an open field**

(A) Laser stimulation protocol for P-DMS animals in an open-field session. Laser was turned on for 1.5 seconds at 4-5mW in 15-second intervals over a 20-minute session. A 1-second window was taken around laser onset and offset for calculating potential laser effects.

(B) Example images show pose captured using Deeplabcut and the mathematical transformation for calculating body angle in laser onset and offset windows.

(C) Velocity surrounding laser onset when laser was delivered unilaterally in open field (n=9; Wilcoxon matched pairs rank sum test, Δvelocity: P = 0.7344)

(D) Velocity surrounding laser offset when laser was delivered unilaterally in open field (n=9; Wilcoxon matched pairs rank sum test, Δvelocity: P = 0.2500)

(E) Body angle surrounding laser onset when laser was delivered unilaterally in open field (n=9; Wilcoxon matched pairs rank sum test, Δangle: P = 0.7344)

(F) Body angle surrounding laser offset when laser was delivered unilaterally in open field (n=9; Wilcoxon matched pairs rank sum test, Δangle: P = 0.3008)

(G) Velocity surrounding laser onset when laser was delivered bilaterally in open field (n=9; Wilcoxon matched pairs rank sum test, Δvelocity: P = 0.8203)

(H) Velocity surrounding laser offset when laser was delivered bilaterally in open field (n=9; Wilcoxon matched pairs rank sum test, Δvelocity: *P = 0.0273)

(I) Body angle surrounding laser onset when laser was delivered bilaterally in open field (n=9; Wilcoxon matched pairs rank sum test, Δangle: $P = 0.1641$)

(J) Body angle surrounding laser offset when laser was delivered bilaterally in open field (n=9; Wilcoxon matched pairs rank sum test, Δangle: $P = 0.7344$)

Supplemental Fig.7

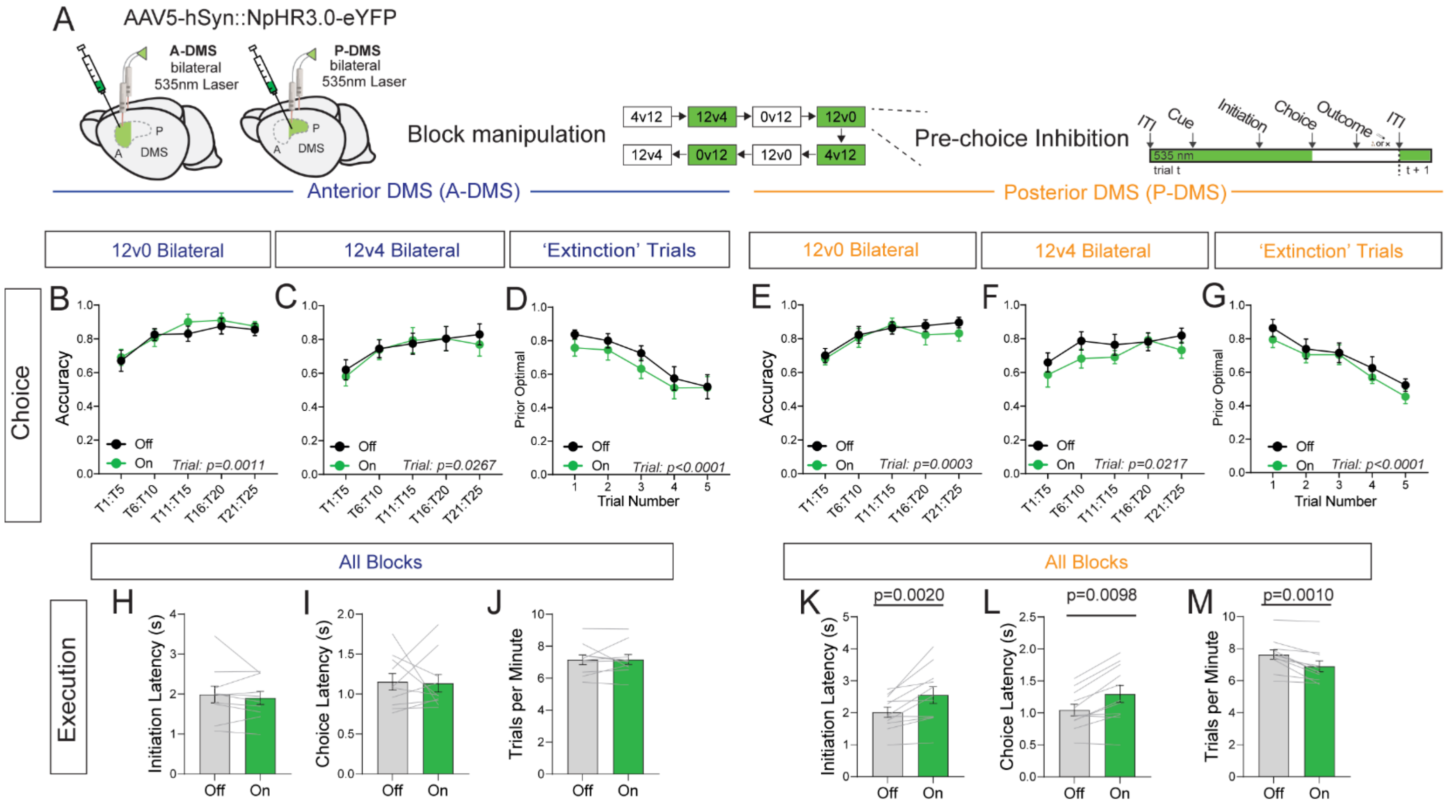


**Supplementary Figure 7. Bilateral pre-choice inhibition slows task execution in P-DMS animals**

(A) Experiment schematic. Bilateral activation of NpHR in the A- or P-DMS in 50% of blocks. Every trial in a manipulation block gets 532nm light delivery during the pre-choice period.

(B) Proportion of optimal choices made by A-DMS:NpHR animals in 12v0 blocks and the laser is off (black) or the laser is on (green) (*n*=10; RM-two-way ANOVA, trial: $F_{2,20} = 9.036$, **P = 0.0011, laser: $F_{1,9} = 0.6019$, P = 0.7003, interaction: $F_{2,20} = 0.3992$, P = 0.7003).

(C) Same as B, but for 12v4 blocks (*n*=10; RM-two-way ANOVA, trial: $F_{2,18} = 4.468$, *P = 0.0267, laser: $F_{1,9} = 0.1261$, P = 0.7307, interaction: $F_{2,19} = 0.2308$, P = 0.8102).

(D) Proportion of choices made by A-DMS:NpHR animals to the prior optimal side in 'extinction' trials when the laser is on (green) or off (black) (*n*=10; RM-two-way ANOVA, trial: $F_{3,24} = 22.36$, ****P < 0.0001, laser: $F_{1,9} = 0.7204$, P = 0.4180, interaction: $F_{3,25} = 0.2754$, P = 0.8290).

(E) Proportion of optimal choices made by P-DMS:NpHR animals in 12v0 blocks and the laser is off (black) or the laser is on (green) (*n*=11; RM-two-way ANOVA, trial: $F_{2,24} = 10.77$, ***P = 0.0003, laser: $F_{1,10} = 0.3449$, P = 0.5700, interaction: $F_{3,28} = 0.5492$, P = 0.6421).

(F) Same as E, but for 12v4 blocks (*n*=11; RM-two-way ANOVA, trial: $F_{3,26} = 4.027$, *P = 0.0217, laser: $F_{1,10} = 2.017$, P = 0.1860, interaction: $F_{2,22} = 0.4901$, P = 0.6329).

(G) Proportion of choices made by P-DMS:NpHR animals to the prior optimal side in 'extinction' trials when the laser is on (green) or off (black) (*n*=11; RM-two-way ANOVA, trial: $F_{3,28} = 16.91$, ****P < 0.0001, laser: $F_{1,10} = 0.7225$, P = 0.4152, interaction: $F_{3,30} = 0.1445$, P = 0.9335).

(H) Initiation latencies for A-DMS:NpHR animals making choices when the laser is off (gray) or on (green) (*n*=10; Wilcoxon matched pairs rank sum test, ΔInit. Latency: P = 0.6250).

(I) Choice latencies for A-DMS:NpHR animals making choices when the laser is off (gray) or on (green) (*n*=10; Wilcoxon matched pairs rank sum test, ∆Choice. Latency: $P = 0.9219$).

(J) Average number of trials completed per minute for A-DMS:NpHR animals when the laser was off (gray) or on (green) (*n*=11; Wilcoxon matched pairs rank sum test, ∆Trial Rate: $P > 0.9999$).

(K) Same as (H) but for P-DMS:NpHR animals (*n*=11; Wilcoxon matched pairs rank sum test, ∆Init. Latency: **$P = 0.0020$).

(L) Same as (I) but for P-DMS:NpHR animals (*n*=11; Wilcoxon matched pairs rank sum test, ∆Choice. Latency: **$P = 0.0098$).

(M) Same as (J) but for P-DMS:NpHR animals (*n*=11; Wilcoxon matched pairs rank sum test, ∆Trial Rate: ***$P = 0.0010$).

Supplemental Fig.8

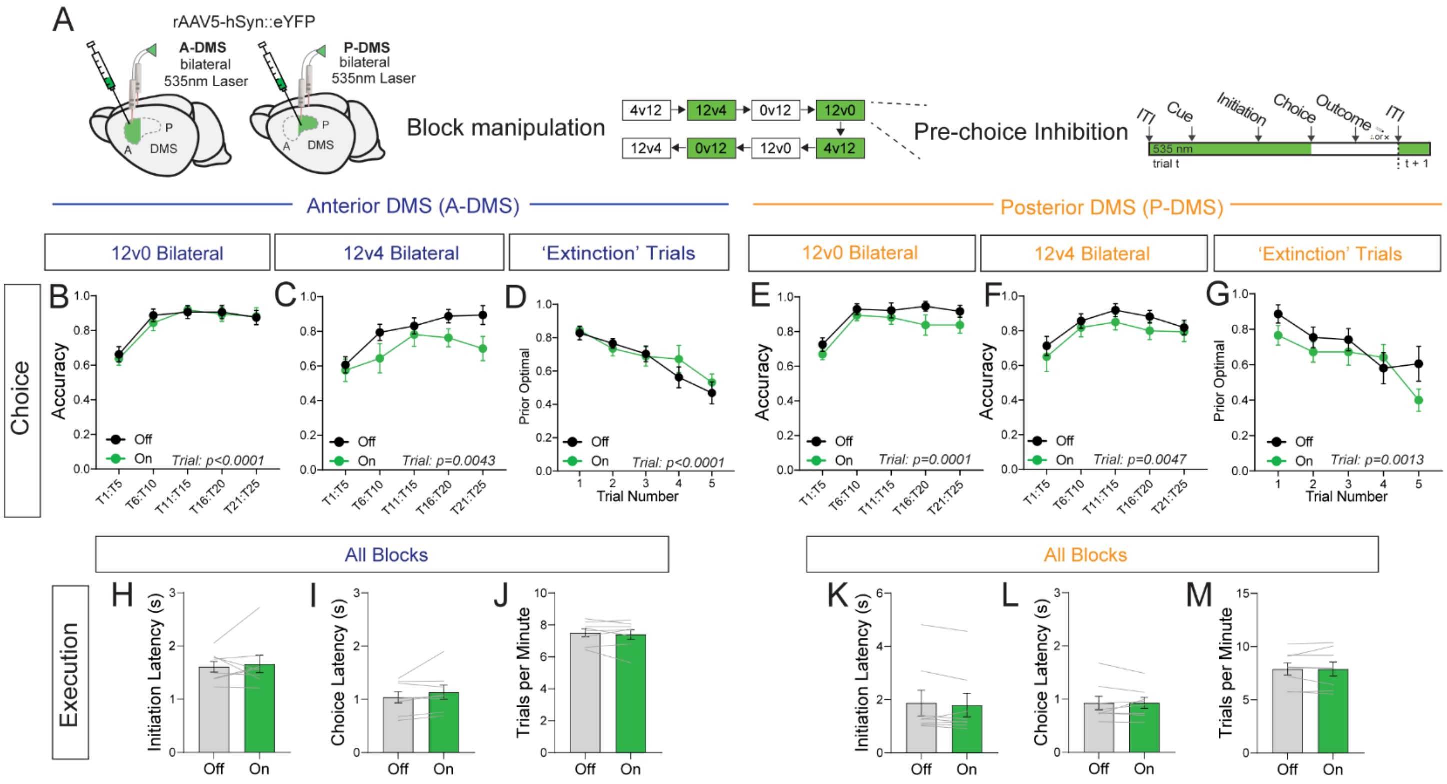


**Supplementary Figure 8. No effects of laser in GFP controls for bilateral pre-choice inhibition**

(A) Experiment schematic. Bilateral activation of GFP in the A- or P-DMS in 50% of blocks. Every trial in a manipulation block gets 532nm light delivery during the pre-choice period.

(B) Proportion of optimal choices made by A-DMS:GFP animals in 12v0 blocks and the laser is off (black) or the laser is on (green) (*n*=8; RM-two-way ANOVA, trial: $F_{2,16}$ = 16.26, ****P < 0.0001, laser: $F_{1,7}$ = 0.8794, P = 0.3796, interaction: $F_{2,17}$ = 0.2284, P = 0.8399).

(C) Same as B, but for 12v4 blocks (*n*=10; RM-two-way ANOVA, trial: $F_{2,13}$ = 8.753, **P = 0.0043, laser: $F_{1,7}$ = 2.921, P = 0.1312, interaction: $F_{2,13}$ = 1.456, P = 0.2672).

(D) Proportion of choices made by A-DMS:GFP animals to the prior optimal side in 'extinction' trials when the laser is on (green) or off (black) (*n*=8; RM-two-way ANOVA, trial: $F_{2,17}$ = 25.05, ****P < 0.0001, laser: $F_{1,7}$ = 0.2717, P = 0.6183, interaction: $F_{2,11}$ = 0.7057, P = 0.4819).

(E) Proportion of optimal choices made by P-DMS:GFP animals in 12v0 blocks and the laser is off (black) or the laser is on (green) (*n*=8; RM-two-way ANOVA, trial: $F_{2,17}$ = 14.96, ***P = 0.0001, laser: $F_{1,7}$ = 3.523, P = 0.1026, interaction: $F_{2,16}$ = 0.3357, P = 0.7525).

(F) Same as E, but for 12v4 blocks (*n*=8; RM-two-way ANOVA, trial: $F_{2,11}$ = 10.53, **P = 0.0047, laser: $F_{1,7}$ = 1.307, P = 0.2905, interaction: $F_{2,16}$ = 0.2288, P = 0.8236).

(G) Proportion of choices made by P-DMS:GFP animals to the prior optimal side in 'extinction' trials when the laser is on (green) or off (black) (*n*=8; RM-two-way ANOVA, trial: $F_{2,17}$ = 9.134, **P = 0.0013, laser: $F_{1,9}$ = 1.484, P = 0.2626, interaction: $F_{2,17}$ = 1.680, P = 0.2140).

(H) Initiation latencies for A-DMS:GFP animals making choices when the laser is off (gray) or on (green) (*n*=8; Wilcoxon matched pairs rank sum test, ∆Init. Latency: P = 0.8438).

(I) Choice latencies for A-DMS:GFP animals making choices when the laser is off (gray) or on (green) (*n*=8; Wilcoxon matched pairs rank sum test, ∆Choice. Latency: P = 0.1094).

(J) Average number of trials completed per minute for A-DMS:GFP animals when the laser was off (gray) or on (green) (*n*=8; Wilcoxon matched pairs rank sum test, ∆Trial Rate: P = 0.7422).

(K) Same as (H) but for P-DMS:GFP animals (*n*=8; Wilcoxon matched pairs rank sum test, ∆Init. Latency: P = 0.2500).

(L) Same as (I) but for P-DMS:GFP animals (*n*=8; Wilcoxon matched pairs rank sum test, ∆Choice. Latency: P = 0.8438).

(M) Same as (J) but for P-DMS:GFP animals (*n*=8; Wilcoxon matched pairs rank sum test, ∆Trial Rate: P = 0.9453).

Supplemental Fig.9

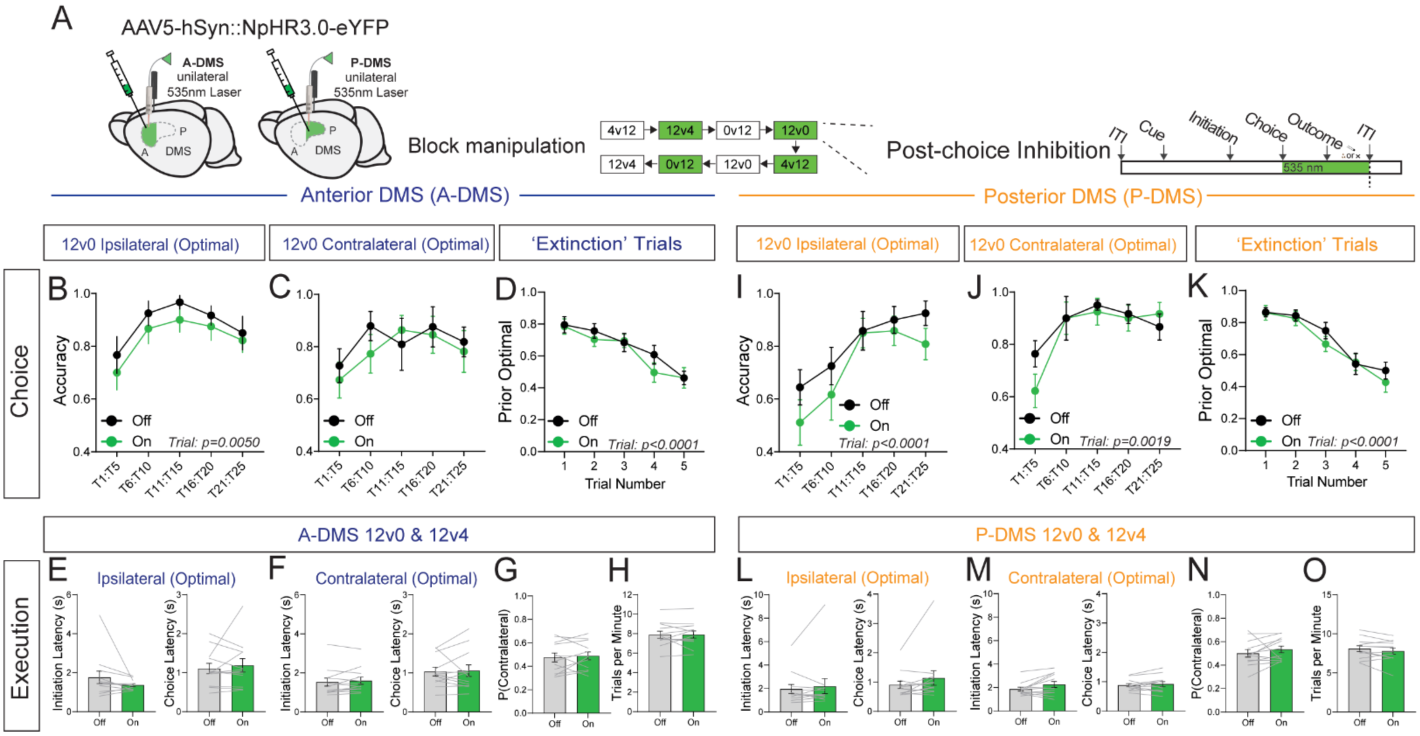


**Supplementary Figure 9. Unilateral post-choice inhibition in 12v0 blocks has no effect on choice or motor execution**

(A) Experiment schematic. Unilateral activation of NpHR in the A- or P-DMS in 50% of blocks. Every trial in a manipulation block gets 532nm light delivery during the post-choice period.

(B) Proportion of optimal choices made by A-DMS:NpHR animals in 12v0 blocks when the laser is ipsilateral to the optimal choice and laser is off (black) or the laser is on (green) (*n*=12; RM-two-way ANOVA, trial: $F_{2,23}$ = 2.528, P = 0.0951, laser: $F_{1,10}$ = 0.9011, P = 0.3649, interaction: $F_{2,24}$ = 0.7759, P = 0.4951).

(C) Same as B, but for trials when the laser is contralateral to the optimal choice (*n*=12; RM-two-way ANOVA, trial: $F_{3,29}$ = 3.448, *P = 0.0309, laser: $F_{1,10}$ = 0.9011, P = 0.3649, interaction: $F_{3,26}$ = 0.7757, P = 0.5019).

(D) Proportion of choices made by A-DMS:NpHR to the prior optimal side in 'extinction' trials when the laser is on (green) or off (black) (*n*=12; RM-two-way ANOVA, trial: $F_{3,30}$ = 19.87, ****P < 0.0001, laser: $F_{1,11}$ = 0.7060, P = 0.4187, interaction: $F_{2,27}$ = 0.9310, P = 0.4233).

(E) Initiation and choice latencies (left and right, respectively) for A-DMS:NpHR animals making choices ipsilateral to the laser manipulation side when the laser was off (gray) or on (green) in the previous trial (*n*=12; Wilcoxon matched pairs rank sum test, Initiation Latency: P = 0.1294, Choice Latency: P = 0.9697).

(F) Same as E, but for trials when the manipulation side is contralateral to the optimal side (*n*=12; Wilcoxon matched pairs rank sum test, Initiation Latency: P = 0.5693, Choice Latency: P = 0.5693).

(G) Probability that A-DMS:NpHR animals made choices to the contralateral side when the laser was off (gray) or on (green) in the previous trial (*n*=12; Wilcoxon matched pairs rank sum test, P(Contralateral): P = 0.6772).

(H) Average number of trials completed per minute for A-DMS:NpHR animals when the laser was off (gray) or on (green) in the previous trial (*n*=12; Wilcoxon matched pairs rank sum test, Trial rate: P = 0.7334).

(I) Proportion of optimal choices made by P-DMS:NpHR animals in 12v0 blocks when the laser is ipsilateral to the optimal choice and laser is off (black) or the laser is on (green) (*n*=12; RM-two-way ANOVA, trial: $F_{2,23}$ = 13.32, ****P = 0.0001, laser: $F_{1,11}$ = 2.217, P = 0.1646, interaction: $F_{2,22}$ = 0.5781, P = 0.5672).

(J) Same as E, but for trials when the laser is contralateral to the optimal choice (*n*=12; RM-two-way ANOVA, trial: $F_{2,23}$ = 8.089, **P = 0.0019, laser: $F_{1,11}$ = 0.3745, P = 0.5530, interaction: $F_{3,34}$ = 1.214, P = 0.3199).

(K) Proportion of choices made by P-DMS:NpHR animals to the prior optimal side in 'extinction' trials when the laser is on (green) or off (black) (*n*=12; RM-two-way ANOVA, trial: $F_{3,29}$ = 8.089, ****P < 0.0001, laser: $F_{1,11}$ = 0.5350, P = 0.4798, interaction: $F_{3,34}$ = 0.5128, P = 0.6509).

(L) Initiation and choice latencies (left and right, respectively) for P-DMS:NpHR animals making optimal choices ipsilateral to the laser manipulation side when the laser was off (gray) or on (green) (*n*=12; Wilcoxon matched pairs rank sum test, Initiation Latency: P = 0.6772, Choice Latency: P = 0.0771).

(M) Same as in L, but for animals making optimal choices contralateral to the laser manipulation side (*n*=12; Wilcoxon matched pairs rank sum test, Initiation Latency: P > 0.0923, Choice Latency: P = 0.5693).

(N) Probability that P-DMS:NpHR animals made choices to the contralateral side when the laser was off (gray) or on (green) (*n*=12; Wilcoxon matched pairs rank sum test, P(Contralateral): P = 0.0771).

(O) Average number of trials completed per minute for P-DMS:NpHR animals when the laser was off (gray) or on (green) (*n*=12; Wilcoxon matched pairs rank sum test, Trial Rate: P = 0.1763).

Supplemental Fig.10

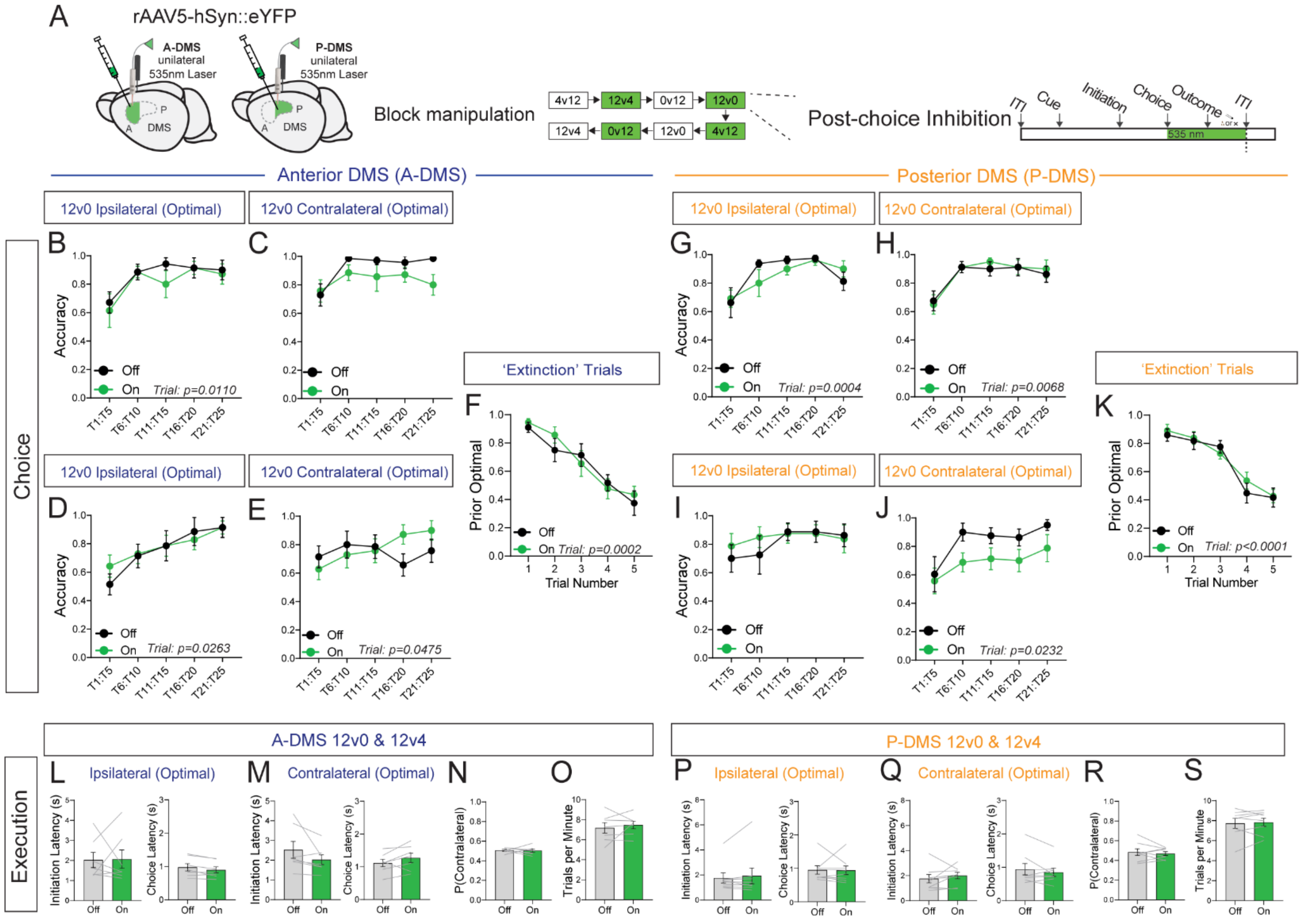


**Supplementary Figure 10. No effects of laser in GFP controls for unilateral post-choice inhibition**
(A) Experiment schematic. Unilateral illumination of GFP in the A- or P-DMS in 50% of blocks. Every trial in a manipulation block gets 532nm light delivery during the post-choice period.

(B) Proportion of optimal choices made by A-DMS:GFP animals in 12v0 blocks when the laser is ipsilateral to the optimal choice and the laser is off (black) or the laser is on (green) (*n*=7; RM-two-way ANOVA, trial: $F_{2,13}$ = 6.266, *P = 0.0110, laser: $F_{1,6}$ = 0.8658, P = 0.3880, interaction: $F_{2,25}$ = 0.3682, P = 0.7394).

(C) Same as B, but for 12v0 blocks when the laser is contralateral to the optimal choice (n=7; RM-two-way ANOVA, trial: $F_{2,13}$ = 2.882, P = 0.0888, laser: $F_{1,7}$ = 5.785, P = 0.0529, interaction: $F_{2,23}$ = 1.402, P = 0.2786).

(D) Proportion of optimal choices made by A-DMS:GFP animals in 12v4 blocks when the laser is ipsilateral to the optimal choice and the laser is off (black) or the laser is on (green) (*n*=7; RM-two-way ANOVA, trial: $F_{2,10}$ = 5.735, *P = 0.0263, laser: $F_{1,7}$ = 0.09499, P = 0.7684, interaction: $F_{3,20}$ = 0.6431, P = 0.5449)

(E) Same as D, but for 12v4 blocks when the laser is contralateral to the optimal choice (*n*=7; RM-two-way ANOVA, trial: $F_{3,16}$ = 3.358, *P = 0.0475, laser: $F_{1,7}$ = 0.09355, P = 0.7700, interaction: $F_{3,20}$ = 1.755, P = 0.2172)

(F) Proportion of choices made by A-DMS:GFP to the prior optimal side in 'extinction' trials when the laser is on (green) or off (black) (*n*=7; RM-two-way ANOVA, trial: $F_{2,12}$ = 20.60, ***P = 0.0002, laser: $F_{1,6}$ = 0.1339, P = 0.7270, interaction: $F_{2,14}$ = 1.136, P = 0.3574)

(G) Proportion of optimal choices made by P-DMS:GFP animals in 12v0 blocks when the laser is ipsilateral to the optimal choice and the laser is off (black) or the laser is on (green) (*n*=8; RM-two-way ANOVA, trial: $F_{3,18}$ = 10.78 ***P = 0.0004, laser: $F_{1,7}$ = 0.1468, P = 0.7130, interaction: $F_{2,13}$ = 1.029, P = 0.3788)

(H) Same as G, but for 12v0 blocks when the laser is contralateral to the optimal choice (*n*=8; RM-two-way ANOVA, trial: $F_{2,14}$ = 7.375, **P = 0.0068, laser: $F_{1,7}$ = 0.2434, P = 0.6369, interaction: $F_{2,12}$ = 0.1623, P = 0.8268)

(I) Proportion of optimal choices made by P-DMS:GFP animals in 12v4 blocks when the laser is ipsilateral to the optimal choice and the laser is off (black) or the laser is on (green) (*n*=8; RM-two-way ANOVA, trial: $F_{1,10}$ = 1.805, P = 0.2148, laser: $F_{1,7}$ = 0.2241, P = 0.6503, interaction: $F_{2,14}$ = 0.6813, P = 0.5257)

(J) Same as H, but for 12v4 blocks when the laser is contralateral to the optimal choice (n=8; RM-two-way ANOVA, trial: $F_{2,13}$ = 5.378, *P = 0.0232, laser: $F_{1,7}$ = 5.485, P = 0.0517, interaction: $F_{3,20}$ = 0.4662, P = 0.7030)

(K) Proportion of choices made by P-DMS:GFP animals to the prior optimal side in 'extinction' trials when the laser is on (green) or off (black) (*n*=8; RM-two-way ANOVA, trial: $F_{3,19}$ = 42.02, ****P < 0.0001, laser: $F_{1,7}$ = 0.1788, P = 0.6851, interaction: $F_{2,11}$ = 0.4107, P = 0.6295)

(L) Initiation and choice latencies (left and right, respectively) for A-DMS:GFP animals making choices ipsilateral to the laser manipulation side when the laser was off (gray) or on (green) (*n*=8; Wilcoxon matched pairs rank sum test, Initiation Latency: P = 0.8125, Choice Latency: P = 0.0781).

(M) Same as in L, but for choices made when the optimal side was contralateral to the laser manipulation side (n=8; Wilcoxon matched pairs rank sum test, Initiation Latency: P = 0.0781, Choice Latency: P = 0.1562).

(N) Probability that A-DMS:GFP animals made choices to the contralateral side when the laser was off (gray) or on (green) (*n*=8; Wilcoxon matched pairs rank sum test, P(Contralateral): P = 0.9375).

(O) Average number of trials completed per minute for A-DMS:GFP animals when the laser was off (gray) or on (green) (*n*=8; Wilcoxon matched pairs rank sum test, Trial Rate: P = 0.5781).

(P) Initiation and choice latencies (left and right, respectively) for P-DMS:GFP animals making choices ipsilateral to the laser manipulation side when the laser was off (gray) or on (green) (*n*=8; Wilcoxon matched pairs rank sum test, Initiation Latency: P > 0.9999, Choice Latency: P = 0.5469).

(Q) Same as P, but for choices made when the optimal side was contralateral to the laser manipulation side (*n*=8; Wilcoxon matched pairs rank sum test, Initiation Latency: P = 0.1953, Choice Latency: P = 0.6406).

(R) Probability that P-DMS:GFP animals made choices to the contralateral side when the laser was off (gray) or on (green) (*n*=8; Wilcoxon matched pairs rank sum test, P(Contralateral): P = 0.5469).

(S) Average number of trials completed per minute for P-DMS:GFP animals when the laser was off (gray) or on (green) (*n*=8; Wilcoxon matched pairs rank sum test, Trial Rate: P = 0.9999).

Supplemental Fig.11

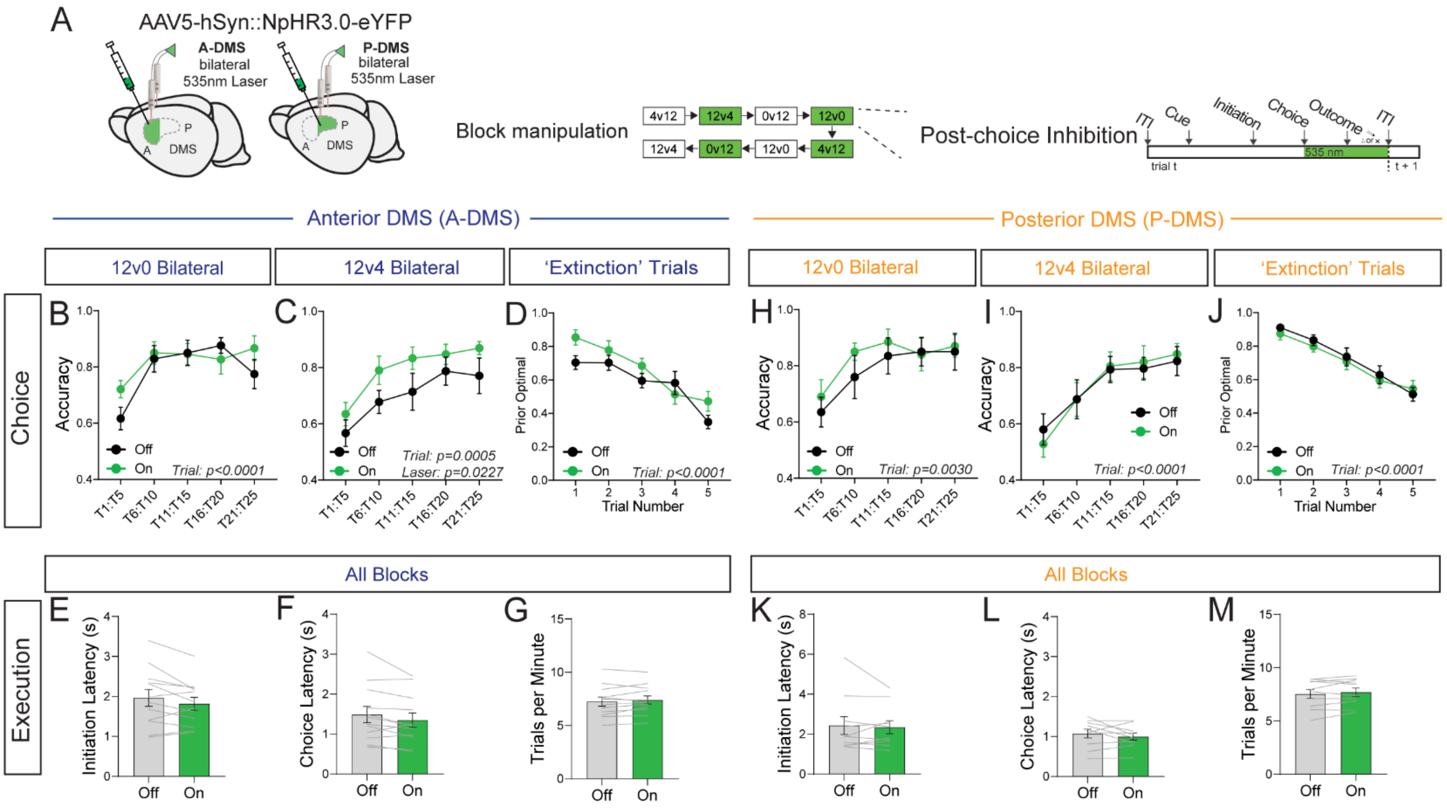


**Supplementary Figure 11. Bilateral post-choice inhibition in A-DMS increases optimal choice selection**
(A) Experiment schematic. Bilateral activation of NpHR in the A- or P-DMS in 50% of blocks. Every trial in a manipulation block gets 532nm light delivery during the post-choice period.

(B) Proportion of optimal choices made by A-DMS:NpHR animals in 12v0 blocks when the laser is off (black) or the laser is on (green) (*n*=12; RM-two-way ANOVA, trial: $F_{3,34}$ = 11.17, ****P < 0.0001, laser: $F_{1,11}$ = 1.195, P = 0.2976, interaction: $F_{2,27}$ = 1.386, P = 0.2688).

(C) Same as B, but for 12v4 blocks (*n*=12; RM-two-way ANOVA, trial: $F_{2,24}$ = 10.31, ***P = 0.0005, laser: $F_{1,11}$ = 7.013, *P = 0.0227, interaction: $F_{2,23}$ = 0.2453, P = 0.7974).

(D) Proportion of choices made by A-DMS:NpHR animals to the prior optimal side in 'extinction' trials when the laser is on (green) or off (black) (*n*=12; RM-two-way ANOVA, trial: $F_{3,24}$ = 22.36, ****P < 0.0001, laser: $F_{1,11}$ = 0.0976, P = 0.0976, interaction: $F_{2,27}$ = 2.003, P = 0.1465).

(E) Initiation latencies for A-DMS:NpHR animals when the laser is off (gray) or on (green) (*n*=12; Wilcoxon matched pairs rank sum test, ΔInit. Latency: P = 0.1099).

(F) Choice latencies for A-DMS:NpHR animals when the laser is off (gray) or on (green) (*n*=12; Wilcoxon matched pairs rank sum test, ΔChoice. Latency: P = 0.0522).

(G) Average number of trials completed per minute for A-DMS:NpHR animals when the laser was off (gray) or on (green) (*n*=12; Wilcoxon matched pairs rank sum test, ΔTrial Rate: P = 0.3013).

(H) Proportion of optimal choices made by P-DMS:NpHR animals in 12v0 blocks and the laser is off (black) or the laser is on (green) (*n*=10; RM-two-way ANOVA, trial: $F_{2,19}$ = 7.667, **P = 0.0030, laser: $F_{1,9}$ = 0.9631, P = 0.3521, interaction: $F_{2,19}$ = 0.2907, P = 0.7591).

(I) Same as H, but for 12v4 blocks (*n*=10; RM-two-way ANOVA, trial: $F_{2,19}$ = 15.49, ****P < 0.0001, laser: $F_{1,9}$ = 0.001007, P = 0.9754, interaction: $F_{3,23}$ = 0.3967, P = 0.7285).

(J) Proportion of choices made by P-DMS:NpHR animals to the prior optimal side in 'extinction' trials when the laser is on (green) or off (black) (*n*=10; RM-two-way ANOVA, trial: $F_{2,22}$ = 48.72, ****P < 0.0001, laser: $F_{1,9}$ = 0.2490, P = 0.6298, interaction: $F_{3,27}$ = 0.3215, P = 0.8132).

(K) Same as E but for P-DMS:NpHR animals (*n*=11; Wilcoxon matched pairs rank sum test, ∆Init. Latency: P = 0.8457).

(L) Same as F but for P-DMS:NpHR animals (*n*=11; Wilcoxon matched pairs rank sum test, ∆Choice. Latency: P = 0.4316).

(M) Same as G but for P-DMS:NpHR animals (*n*=11; Wilcoxon matched pairs rank sum test, ∆Trial Rate: P = 0.4316).

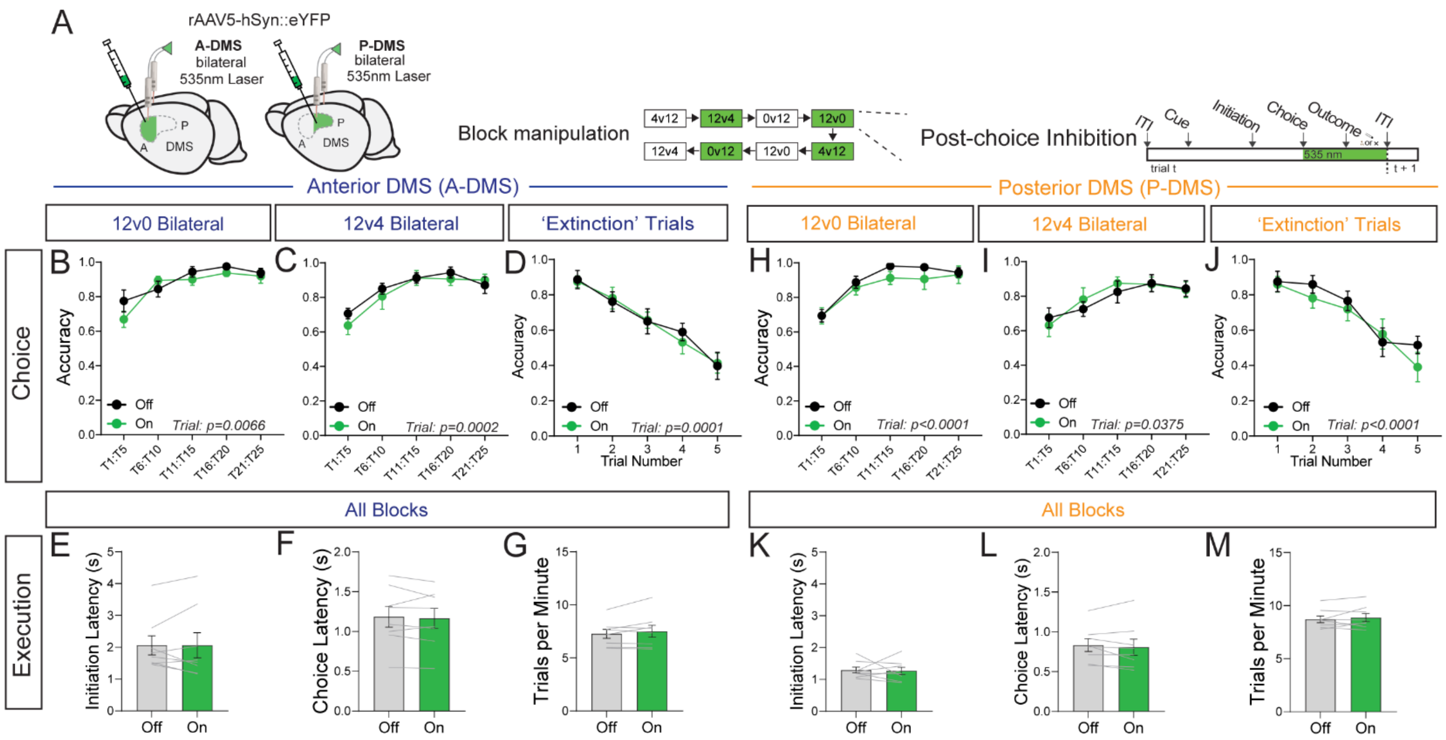

**Supplementary Figure 12. No effects of laser in GFP controls for bilateral post-choice inhibition**
(A) Experiment schematic. Bilateral activation of GFP in the A- or P-DMS in 50% of blocks. Every trial in a manipulation block gets 532nm light delivery during the post-choice period.

(B) Proportion of optimal choices made by A-DMS:GFP animals in 12v0 blocks when the laser is off (black) or the laser is on (green) (*n*=8; RM-two-way ANOVA, trial: $F_{2,12}$ = 8.708, **P < 0.0066, laser: $F_{1,7}$ = 4.055, P = 0.0839, interaction: $F_{3,22}$ = 1.716, P = 0.1923).

(C) Same as B, but for 12v4 blocks (*n*=8; RM-two-way ANOVA, trial: $F_{2,24}$ = 10.31, ***P = 0.0005, laser: $F_{1,11}$ = 7.013, *P = 0.0227, interaction: $F_{2,23}$ = 0.2453, P = 0.7974).

(D) Proportion of choices made by A-DMS:GFP animals to the prior optimal side in 'extinction' trials when the laser is on (green) or off (black) (*n*=8; RM-two-way ANOVA, trial: $F_{2,15}$ = 16.71, ***P = 0.0001, laser: $F_{1,7}$ = 0.01955, P = 0.8927, interaction: $F_{3,21}$ = 0.1873, P = 0.9024).

(E) Initiation latencies for A-DMS:GFP animals when the laser is off (gray) or on (green) (*n*=8; Wilcoxon matched pairs rank sum test, ∆Init. Latency: P = 0.9453).

(F) Choice latencies for A-DMS:GFP animals when the laser is off (gray) or on (green) (*n*=8; Wilcoxon matched pairs rank sum test, ∆Choice. Latency: P = 0.3828).

(G) Average number of trials completed per minute for A-DMS:GFP animals when the laser was off (gray) or on (green) (*n*=8; Wilcoxon matched pairs rank sum test, ∆Trial Rate: P = 0.7422).

(H) Proportion of optimal choices made by P-DMS:GFP animals in 12v0 blocks and the laser is off (black) or the laser is on (green) (*n*=8; RM-two-way ANOVA, trial: $F_{2,13}$ = 38.48, ****P < 0.0001, laser: $F_{1,7}$ = 1.037, P = 0.3425, interaction: $F_{2,16}$ = 0.3246, P = 0.7559).

(I) Same as H, but for 12v4 blocks (*n*=8; RM-two-way ANOVA, trial: $F_{2,11}$ = 4.907, *P = 0.0375, laser: $F_{1,7}$ = 0.1315, P = 0.7276, interaction: $F_{2,15}$ = 0.8698, P = 0.4443).

(J) Proportion of choices made by P-DMS:GFP animals to the prior optimal side in ‘extinction’ trials when the laser is on (green) or off (black) (*n*=8; RM-two-way ANOVA, trial: $F_{3,20}$ = 22.54, ****P < 0.0001, laser: $F_{1,7}$ = 0.4.133, P = 0.0816, interaction: $F_{3,17}$ = 0.6258, P = 0.5736).

(K) Same as E but for P-DMS:GFP animals (*n*=8; Wilcoxon matched pairs rank sum test, ∆Init. Latency: P = 0.8438).

(L) Same as F but for P-DMS:GFP animals (*n*=8; Wilcoxon matched pairs rank sum test, ∆Choice Latency: P = 0.3828).

(M) Same as G but for P-DMS:GFP animals (*n*=8; Wilcoxon matched pairs rank sum test, ∆Trial Rate: P = 0.6406).

Supplemental Fig.13

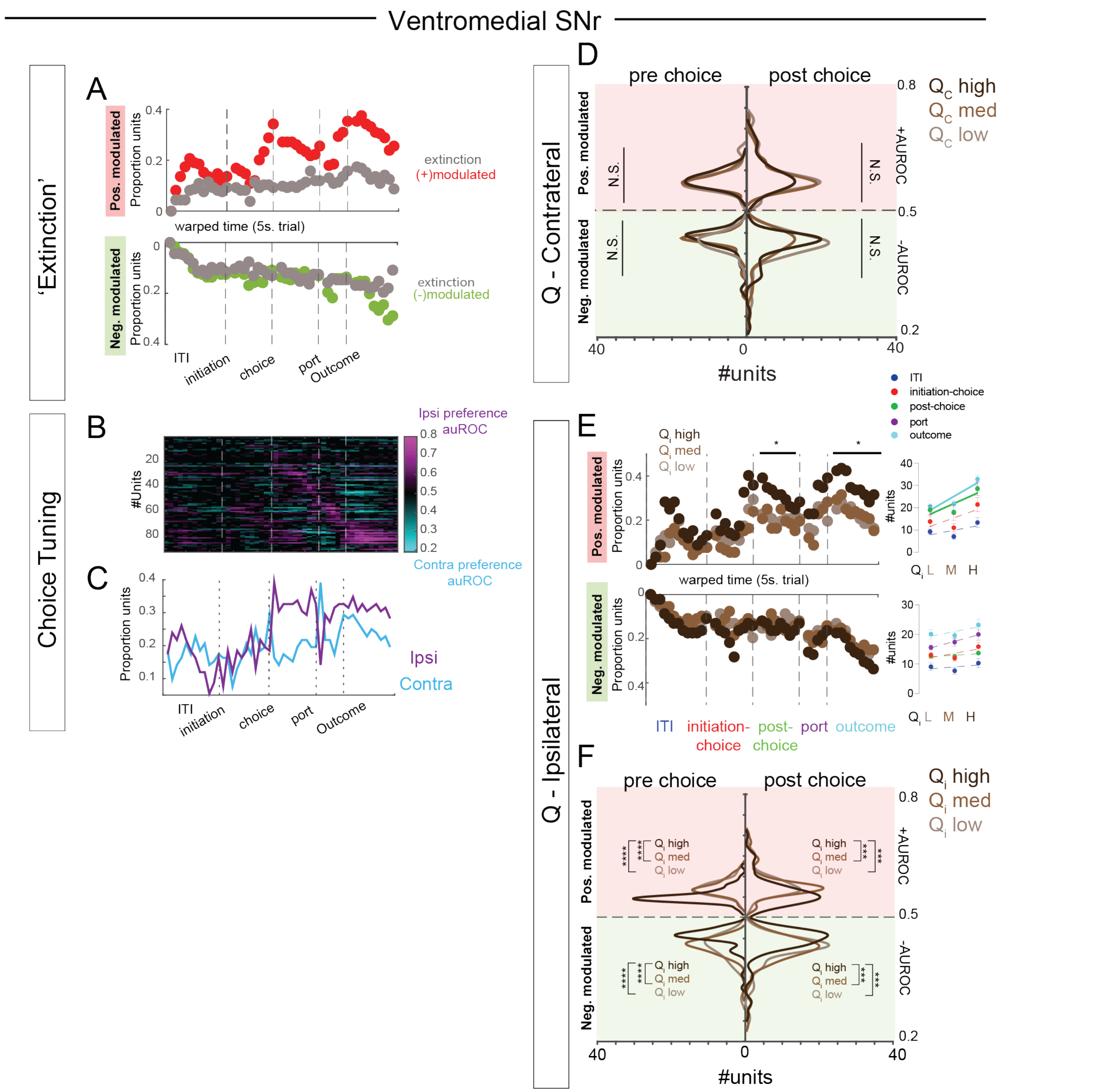


**Supplementary Figure 13. Changes in SNr activity for extinction trials, choice direction, and value**

(A) Top: reduced number of positively modulated units in the SNr as animals during extinction trials compared to non-extinction trials. Bottom: No changes in negative modulation for extinction trials compared to non-extinction trials.

(B) Individual unit choice tuning for SNr neurons as measured by the presence of increased or reduced auROC scores in ipsilateral vs contralateral choices.

(C) Proportion of SNr neurons with ipsilateral or contralateral choice tuning throughout a trial.

(D) Plot of the number of modulated units (x-axis) and their respective modulation indices (auROC, y-axis) for three $Q_{contra}$ values, separated by positive (pink) and negative (green) modulation as well as by pre- or post-choice (left versus right half). Individual SNr units did not scale activity towards higher auROC values as a function of value for positive modulation in the pre-choice period (Top Left: Kruskal-Wallis test with Dunn's test for multiple comparisons, $H_2$ = 2.002, P=0.3676) or the post-choice period (Top Right: Kruskal-Wallis test with Dunn's test, $H_2$ = 0.2705, P=0.8735). Similarly, units did not scale their activity towards lower auROC values as a function of value for negative modulation in the pre-choice period (Bottom Left: Kruskal-Wallis test, $H_2$ = 2.886, P=0.2362) or the post-choice period (Bottom Right: Kruskal-Wallis test, $H_2$ = 0.4085, P=0.8152).

(E) Top: changes in the proportion of positively modulated units at different values for ipsilateral choice in the SNr. Positive modulation scales positively with value in the post-choice window and the outcome window for the SNr. Bottom: Negative modulation does not with ipsilateral choice value in the SNr. Insets show linear regression for number of units modulated within each window according to value.

(F) Same as D, but for $Q_{ipsi}$ values. auROC values for individual units reveal that there is increased positive modulation of SNr units for lower $Q_{ipsi}$ values in both the pre-choice period (Top Left: Kruskal-Wallis test with Dunn's test for multiple comparisons, $H_2$ = 49.69, ****P<0.0001; *Post-hoc* comparisons: ****$p<0.0001$) and the post-choice period (Top Right: Kruskal-Wallis test with Dunn's test for multiple comparisons, $H_2$ = 20.60, ****P<0.0001; *Post-hoc* comparisons: ***$p<0.001$). Individual units also scaled their activity towards increased negative modulation as a function of value in the SNr for the pre-choice period (Bottom Left: Kruskal-Wallis test with Dunn's test for multiple comparisons, $H_2$ = 31.47, ****P<0.0001; *Post-hoc* comparisons: ****p<0.0001) and the post-choice period (Bottom Right: Kruskal-Wallis test with Dunn's test for multiple comparisons, $H_2$ = 19.25, ****P<0.0001; *Post-hoc* comparisons: ***p<0.001).
